# Neuroactive steroids activate membrane progesterone receptors to induce sex specific effects on protein kinase activity

**DOI:** 10.1101/2025.01.24.634751

**Authors:** Abigail H. S. Lemons, Briana Murphy, Jake S. Dengler, Seda Salar, Paul A. Davies, Joshua L. Smalley, Stephen J. Moss

## Abstract

Neuroactive steroids (NAS), which are synthesized in the brain from progesterone, exert potent effects on behavior and are used to treat postpartum depression, yet how these compounds induce sustained modifications in neuronal activity are ill-defined. Here, we examined the efficacy of NAS for membrane progesterone receptors (mPRs) δ and ε, members of a family of GPCRs for progestins that are expressed in the CNS. NAS increase PKC activity via G_q_ activation of mPRδ with EC50s between 3-11nM. In contrast, they activate G_s_ via mPRε to potentiate PKA activity with similar potencies. NAS also induced rapid internalization of only mPRδ. In the forebrain of female mice, mPRδ expression levels were 8-fold higher than males. Consistent with this, activation of PKC by NAS was evident in acute brain slices from female mice. Collectively, our results suggests that NAS may exert sex-specific effects on intracellular signaling in the brain via activation of mPRs.

## Introduction

Neuroactive steroids (NAS) such as allopregnanolone (ALLO) are synthesized in the brain from progesterone (P4) and exert potent anticonvulsant, anxiolytic, and sedative actions.^1,2^ ZULRESSO^®^ a proprietary formulation of brexanolone (ALLO), is used to treat postpartum depression (PPD). ZULRESSO^®^exhibits fast onset within 60 hours and its efficacy is then maintained for at least 30 days following drug withdrawal.^3,4^ Another synthetic NAS, Ganaxalone, is used to arrest refractory seizures in cyclin-dependent kinase-like 5 (CDKL5) deficiency disorder.^5^ Mechanistically, some NAS, like ALLO, exert acute effects on neuronal excitability via their efficacy as positive allosteric modulators (PAMs) of both synaptic and extrasynaptic γ-aminobutyric type A receptors (GABA_A_R), which mediate phasic and tonic inhibition in the adult brain, respectively.^6^ The efficacy of NAS that function as GABA_A_R PAMs is dependent upon a common binding site that is found in all subtypes of this heterogenous family of pentameric ligand-gated ion channels.^7–9^

In addition to ALLO, novel synthetic NAS like SGE-516 have also been used to investigate the effects of NAS in the brain (Figure 1). SGE-516 has similar pharmacological properties to ALLO on GABA_A_Rs and the magnitude of phasic in addition to tonic inhibition. Like ALLO, SGE-516 exerts anxiolytic, anticonvulsant, anti-depressant and sedative efficacies in rodents, but has increase bioavailability.^10–14^

**Figure 1.**
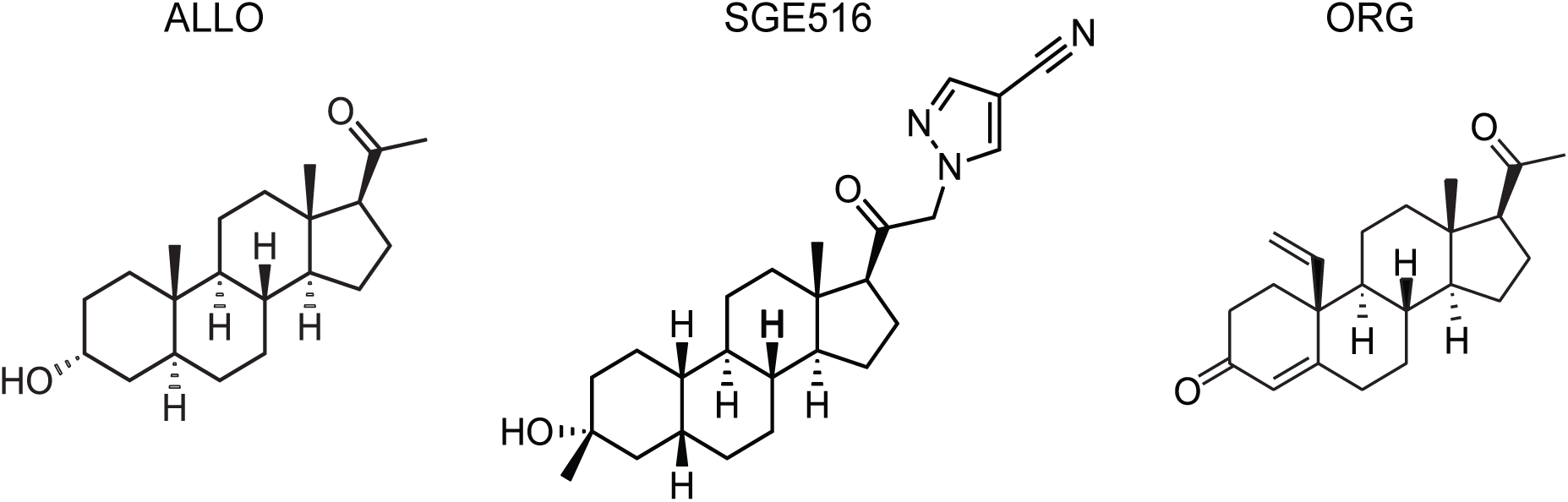
Chemical structures of Allopregnanolone, SGE-516, and Org OD 02-0. The chemical structures of the three drug treatments used to assess the signaling of mPRs. The structure of allopregnanolone (ALLO), SGE-516, and Org OD 02-0 (ORG) are shown from left to right.

Some NAS, including ALLO and SGE-516, also regulate GABA_A_R phosphorylation and insertion into the plasma membrane, a process dependent on cAMP dependent kinase (PKA) and protein kinase C (PKC) effects that are independent of NAS efficacy as GABA_A_R PAMs.^10,15,16^ One possible mechanism to explain these pleiotropic effects of NAS is that they are by metabotropic “NAS” receptors. Consistent with this concept, studies in yeast, fish, mammalian cell lines and reproductive tissues have revealed that progestins can bind to a family of membrane progesterone receptors (mPR) with five members ɑ, β, γ, δ, and ε, that share homology with classical serpentine G-protein coupled receptors (GPCRs).^17–22^. In contrast to this, mPRs have been reported to signal via G-protein independent mechanisms, and their failure to signal in some cell lines has been attributed to intracellular retention.^18,22–24^ The membrane topology of mPRs has been analyze using homology modelling and immunohistochemistry when expressed in cell line. Currently these is no consensus of the number of transmembrane or the localization of the N- and C-terminus of mPRs (see; Thomas 2022 for a review).^18^

Here, we examined the roles that mPRδ and mPRε play in mediating the effects of ALLO and SGE-516 in addition to the mPR agonist ORG-OD-020 (ORG) on intracellular signaling. We focused on mPRδ and mPRε because they are two mPR subtypes predicted to increase intracellular signaling.^25,26^ Our results demonstrate that ALLO activates mPRε leading to PKA activation via G_s_ signaling. In contrast, both ALLO and SGE-516 treatment increased PKC activity that was dependent upon mPRδ via G_q_ signaling suggesting that SGE-516 is a potent agonist for mPRδ. In acute hippocampal brain slices, SGE-516 selectively increased PKC activity only in slices prepared from female mice. Consistent with this, 8-fold higher expression levels of mPRδ mRNA was seen in the brains of adult female mice. Thus, NAS, like SGE-516, may mediate sustained and sex specific effects on neuronal signaling via distinct mPR isoforms.

## Results

### Creation of mPR-expressing stable cell lines to be used as tools to investigate signaling following NAS treatment

To study the role of mPRs in mediating metabotropic signaling by NAS we created clonal MDA-MB-231 (MDA231) cell lines expressing individual mPR isoforms. MDA231 cells, derived from a triple negative breast cancer tumor, do not express any nuclear hormone receptors and express very low levels of only mPRɑ, making these cell lines ideal candidates to explore intracellular signaling following exposure to NAS.^27,28^ Our studies focused on mPRδ and ε, as these isoforms are highly expressed in the brain, and based on homology with other GPCRs, are predicted to activate the stimulatory G-protein G_s_.^18,26^ To directly visualize their expression, the respective proteins were modified at their C-terminus with GFP as our reporter epitope.

To create the stable mPR-expressing cell lines, mPRδ-GFP and mPRε-GFP constructs were transduced into MDA231 cells using lentiviral vectors. GFP positive cells were then selected for mPR-GFP expression using three rounds of Fluorescence Activated Cell Sorting (FACS). During the final sort, individual cells were placed into each well of a 96-well plate. The GFP-fluorescence and growth rate for each of the 96 resulting clonal lines was assessed, and the optimal clone was selected for further use (Supplemental Figure 1). Some studies have suggested that in expression systems mPRs are retained within the endoplasmic reticulum (ER)^19,29^. To examine this in our cell lines, they were immunostained with antibodies against GFP and BAP31, an ER resident protein, followed by confocal imaging.^30–32^ Robust GFP/BAP31 puncta were detected in both cell lines in intracellular compartments, in addition to puncta positive for GFP alone. To quantify these results, we used a colocalization plugin in FIJI to compare the percent of GFP puncta that co-localized with BAP31. The majority of GFP co-localized with BAP31, (∼60-80%; Figure 2B) as membrane proteins are synthesized in the ER and reach conformation maturity in this organelle. However, 21.3±4.3 and 41.5±2.7 of mPRδ-GFP and mPRε-GFP puncta did not co-localize with BAP31 (Figure 2C).

**Figure 2.**
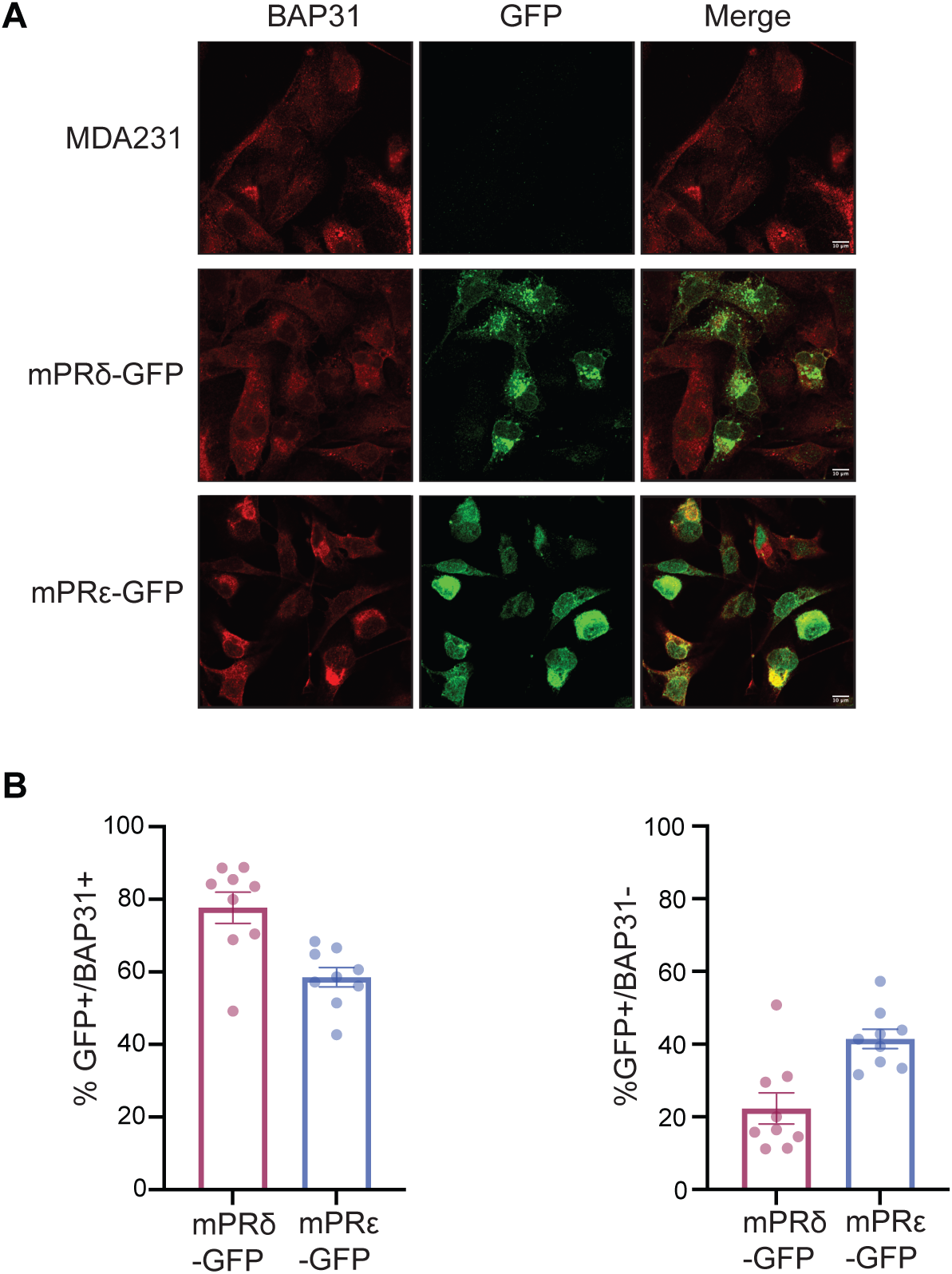
Examining the subcellular distribution of mPRs in MDA231 cells. **A.** Cell lines were fixed and immunostained with BAP31 and GFP antibodies and imaged by confocal microscopy. Scale bar represents 10μm. **B**. The number of GFP puncta that were positive and negative for BAP31 immunoreactivity was then determined using a colocalization plugin in ImageJ/Fiji. (N= 3, *n*=3). In all panels, data represents mean ± SEM.

To further evaluate the subcellular localization, cultures were permeabilized immunostained with actin and GFP antibodies, and subject to Total internal reflection fluorescence microscopy (TIRF). The use of TIRF microscopy allows for measurement of fluorescently-tagged surface protein expression as only protein within 100nm of the cell surface will fluoresce when imaged.^33^. Robust fluorescence was seen for mPRδ-GFP and mPRε-GFP expressing cells but not in untransfected controls (Supplemental Figure 2).

Collectively these results demonstrate that a proportion of both mPRs are ER-transport competent and accumulate in distinct compartments within the secretory or endocytic pathways. The TIRF imaging further demonstrates that mPRδ-GFP and mPRε-GFP accumulate on or within 100nM of the plasma membrane.

### ORG selectively activates PKC and PKA dependent upon mPRδ and mPRε

To gain initial insights into mPR signaling, we examined the effects of ORG on the activity of PKA, PKC, and Src. accepted effectors for Gs, Gq, Gi respectively. Kinase activity was determined using phospho-antibodies that recognize the phosphorylated, active kinase and antibodies against non-phosphorylated epitopes. Kinase activation was measured by phosphorylation at specific amino acid residues: Thr197protein (PKA), Thr514 (PKC), and Tyr416 (Src).^34–39^ The ratios of phosphorylated to total kinase immunoreactivity were determined and normalized to control treatment within each cell line and then normalized to the response in the control MDA231 cells. An incubation time of 20 minutes was chosen based on an initial pilot experiment and prior kinase-dependent biochemical and electrophysiological changes after NAS treatment (Supplemental Figure 3).^10,15,16^

In mPRδ-GFP cells, a 106.69 ± 24.79% and 113.88 ± 25.71% increase compared to control treatment (100%) in PKC activation was observed following 100nM and 300nM ORG treatments, respectively (3nM: 112.40 ± 18.60%, p=0.9970; 10nM: 127.69 ± 8.98%, p=0.8893; 30nM: 146.78 ± 44.93%, p=0.2622; 100nM: 206.69 ± 24.79%, p=0.0024; 300nM: 213.88 ± 25.71%, p<0.0001; Figure 3A-B). No significant effects on PKA (3nM: 102.24 ± 6.13%, p>0.9999; 10nM: 97.651 ± 4.68%, p=0.6482; 30nM: 87.445 ± 4.70%, p>0.9999; 100nM: 97.241 ± 6.11%, p=0.8920; 300nM: 104.36 ± 6.27%, p=9705) or Src (3nM: 103.47 ± 6.03%, p=0.8138; 10nM: 106.54 ± 3.90%, p=0.7083; 30nM: 112.01 ± 7.52%, p=0.1065; 100nM: 92.669 ± 12.01%, p=0.9984; 300nM: 108.98 ± 7.53%, p=0.9944) were seen with ORG in this cell line (Supplemental Figure 4).

**Figure 3.**
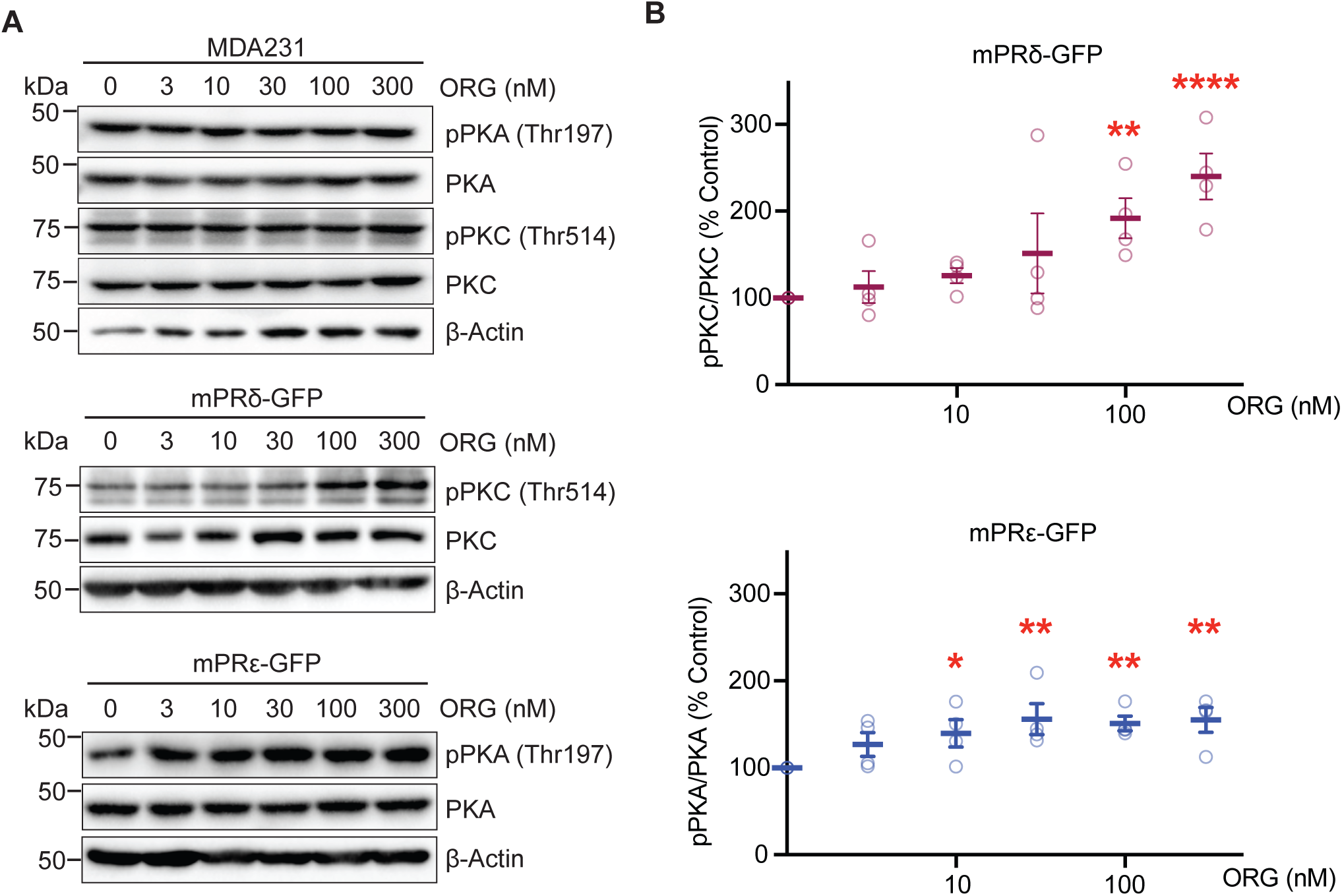
Examining the effects of ORG on PKA and PKC activity in cell lines expressing mPRs. **A.** Representative immunoblots for MDA231, mPRδ-GFP, and mPRε-GFP cells treated for 20 minutes with the pan-mPR agonist, ORG at 3, 10, 30, 100, and 300nM. Cells were lysed, the proteins resolved on SDS-PAGE and subject to immunblotting. **B.** Densitometry was carried out to quantify the difference between host and mPR expressing cells and between ORG treatment concentrations. The ratio of phosphorylated kinase to total kinase was determined and normalized to values in MDA231 cells (100%). Two-Way ANOVA was used to compare treatment responses in each cell line. Post hoc comparisons of each ORG concentration were calculated using the Šidák multiple comparison test. * p<0.05, ** p<0.01, *** p<0.001 (*n*=4). In all panels, data represents mean ± SEM.

For mPRε-expressing cells, a 57.89% increase in PKA activation following 10nM ORG compared to control treatment response (100%) was observed (10nM: 157.89 ± 17.85%, p=0.0136). PKA activation was also observed at higher ORG treatment concentrations (30nM: 156.37 ± 18.00%, p=0.0015; 100nM: 158.75 ± 8.94%, p=0.0040; 300nM: 170.29 ± 15.82% p=0.0017; Figure 3A-B). In contrast to this, ORG did not significantly modify PKC (3nM: 96.618 ± 3.49%, p=0.9997; 10nM: 105.21 ± 8.03%, p=0.997; 30nM: 97.008 ± 10.90%, p>0.9999; 100nM: 100.93 ± 4.46%, p=0.9983; 300nM: 104.73 ± 15.95%, 0.9716), or Src activity (3nM: 99.863 ± 2.71%, p=0.9314; 10nM: 94.089 ± 5.09%, p=0.1820; 30nM: 99.012 ± 10.19%, p=0.5695; 100nM: 112.65 ± 8.58%, p=0.9221; 300nM: 113.27 ± 10.32%, 0.9308; Supplemental Figure 4) in mPRε-GFP cells. Thus, ORG selectively activates PKC or PKA activity dependent upon mPRδ and mPRε, respectively.

### ORG treatment selectively regulates the cell surface stability of mPRδ

To analyze the effects of ORG on the plasma membrane stability of mPRs, total internal reflection fluorescence (TIRF) microscopy was used to perform live imaging of the mPRδ-GFP and mPRε-GFP cells following acute treatment with ORG. The use of TIRF microscopy allows for measurement of fluorescently-tagged surface protein expression as only protein within 100nm of the cell surface will fluoresce when imaged.^33^ Both mPRδ-GFP and mPRε-GFP cells exhibited robust TIRF signal as measured using live imaging after fluorescence levels recovered to baseline following photobleaching due to the process of focusing the laser consistent with their accumulation on or near the plasma membrane (Figure 2, Supplemental Figure 2).

Classical GPCRs undergo rapid agonist induced endocytosis, a process that regulates their lifetime on the plasma membrane, and the specificity of the effectors they activate.^40^ To assess if mPRs undergo this process, mPRδ-GFP and mPRε-GFP cells were subject to TIRF imaging for 20 minutes in the presence of 100nM ORG or vehicle. The TIRF signal was then normalized to that seen for cells exposed to vehicle. For mPRδ-GFP the TIRF signal was reduced by approximately 20% compared to vehicle (p=0.0007; Figure 4B). This reduction is consistent with a reduction of mPRd accumulation on or with 100nM of the plasma membrane. In contrast, no significant change in the TIRF was seen for mPRε-GFP under the same conditions. Collectively, our measurements using TIRF suggest that ORG treatment induces selective internalization of mPRδ receptors.

**Figure 4.**
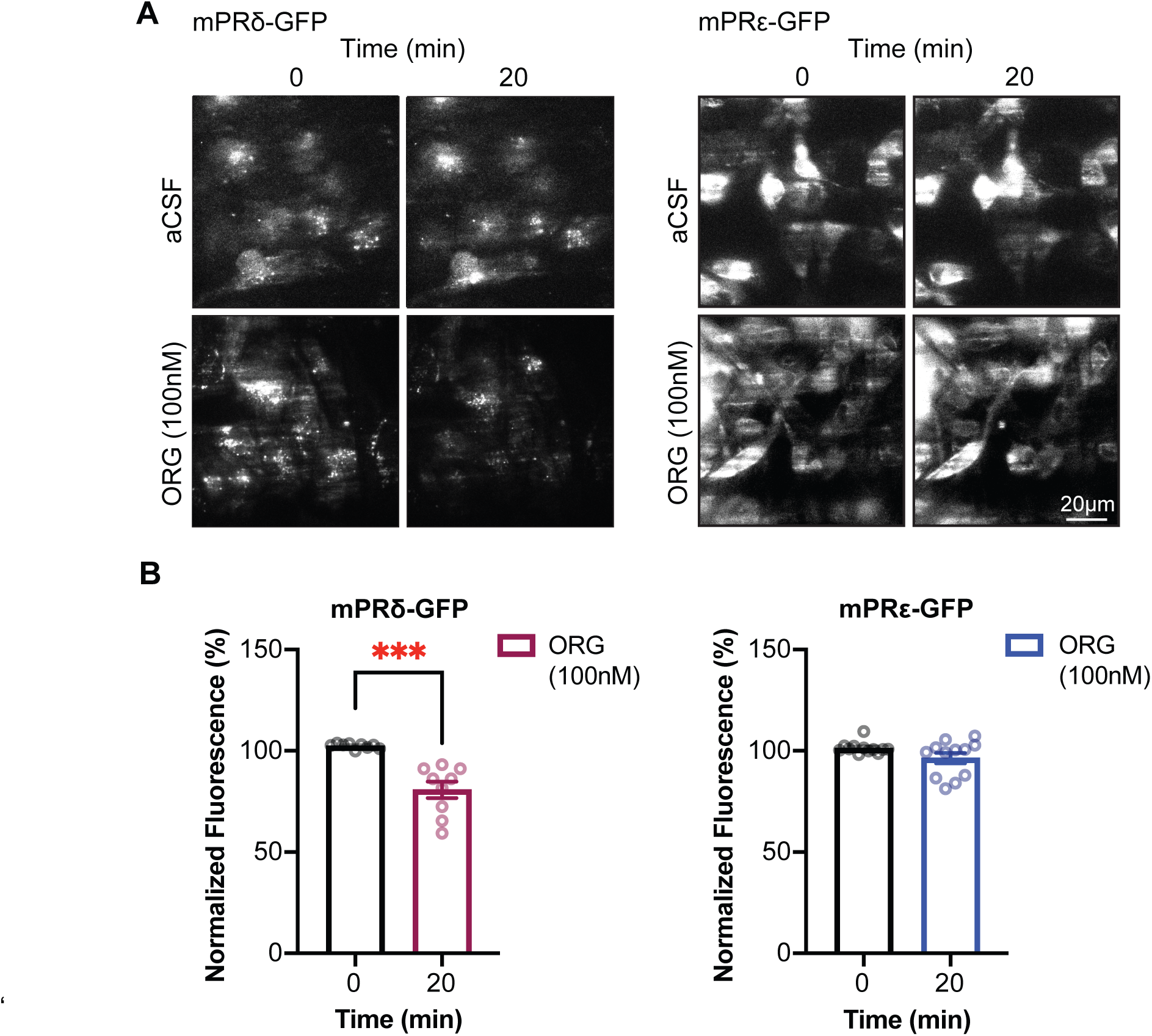
Examining the effects of ORG on mPR cell surface stability. **A**. Representative images of GFP signal in cells imaged using TIRF microscopy following 20-minute treatment with aCSF or 100nM mPR agonist, ORG. **B**. Quantified and normalized fluorescence comparing baseline to ORG treatment for 20 minutes in mPRδ-GFP and mPRε-GFP cells. Data were normalized to aCSF treatment within each cell line. *** p<0.001, N=3, *n*=9-12. In all panels, data represents mean ± SEM.

### ALLO activates PKC and PKA dependent upon mPRδ and mPRε

Next, we investigated whether ALLO, an endogenous NAS, activated mPRδ or mPRε in our cellular system. Immunoblots for total and phosphorylated PKA and PKC revealed that ALLO treatment led to a 43.58 ± 5.34% increase in PKC phosphorylation compared to control treatment in mPRδ−GFP cells that was significant at 300nM (3nM: 84.378 ± 13.74%, p=0.9999; 10nM: 99.569 ± 16.97%, p=0.9468; 30nM: 106.64 ± 10.27%, p=0.4674; 100nM: 115.67 ± 3.82%, p=0.1022; 300nM: 143.58 ± 5.34%, p=0.0004; Figure 5A-B). ALLO did not modify PKA activity at any dose tested in mPRδ-GFP cells (3nM: 86.327 ± 5.39%, p=0.9886; 10nM: 91.441± 4.66%, p>0.9999; 30nM: 83.155 ± 4.54%, p=0.7576; 100nM: 98.624 ± 11.00%, p=0.9949; 300nM: 103.26 ± 14.05%, p>0.9999; Supplemental Figure 5).

**Figure 5.**
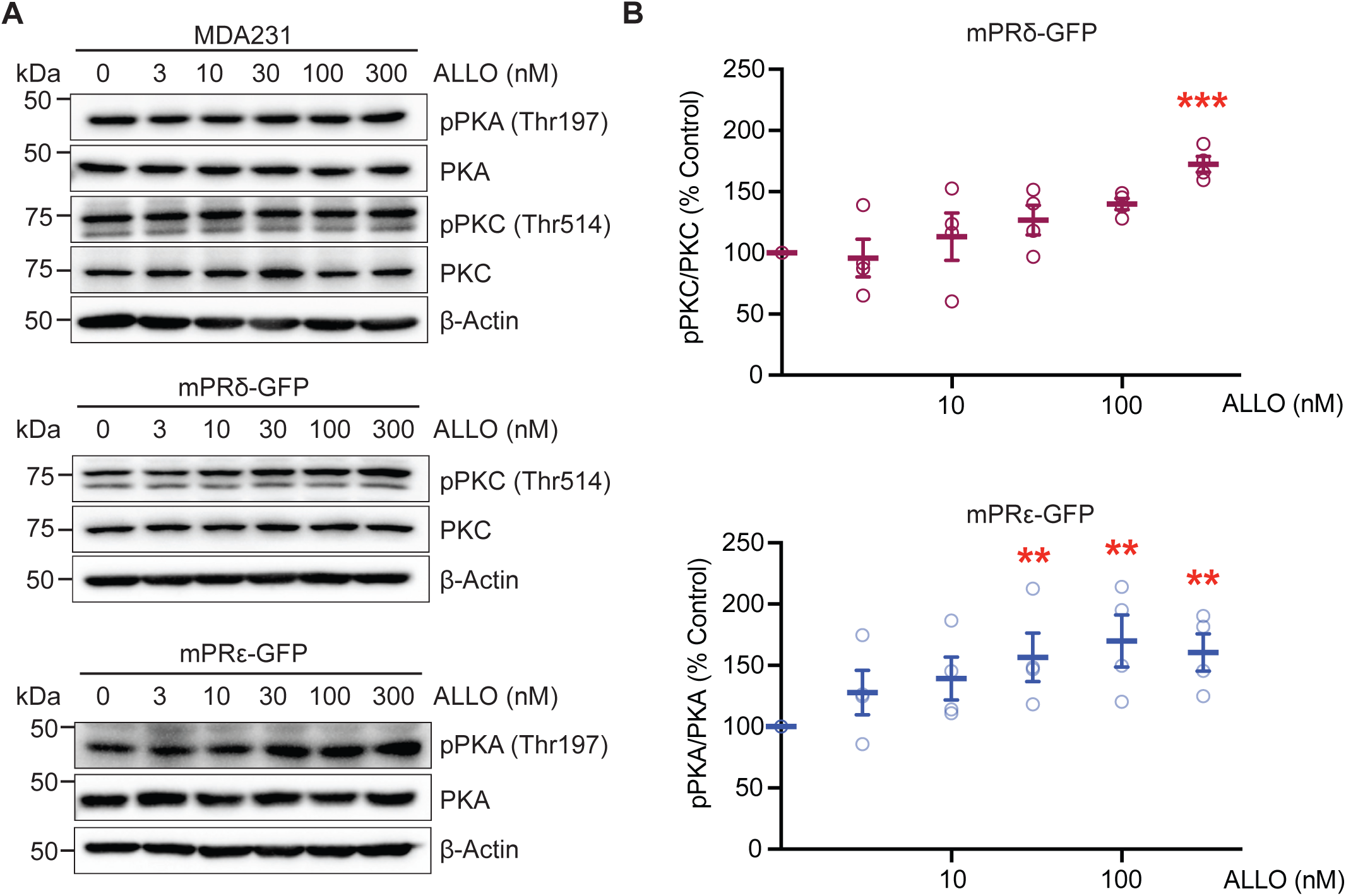
Measuring the effects of ALLO on mPR signaling. **A.** Representative immunoblots for MDA231, mPRδ-GFP, and mPRε-GFP cells treated for 20 minutes with ALLO at 3, 10, 30, 100, and 300nM. Cells were lysed, the proteins resolved on SDS-PAGE and subject to immunblotting**. B**. Densitometry was carried out to quantify the difference between host and mPR expressing cells and between ALLO treatment concentrations. The ratio of phosphorylated kinase to total kinase was determined and normalized to control treatment (100%). Two-Way ANOVA was used to compare treatment responses relative to control MDA231 cells. Post hoc comparisons of each ALLO concentration were calculated using the Šidák multiple comparison test. * p<0.05, ** p<0.01, *** p<0.001, n=4. In all panels, data represents mean ± SEM.

In the mPRε−GFP cells, ALLO treatment increased PKA activation by 53.44% at 30nM (153.4 ± 19.57, p=0.0028), 60.49% at 100nM (160.49 ± 20.14%, p=0.0056), and 64.44% at 300nM (164.44 ± 15.71%, p=0.0046), but had no effect at lower concentrations (3nM: 105.57 ± 15.02%, p=0.3688; 10nM: 130.55 ± 16.45%, p=0.1861; Figure 5A-B). Additionally, no effects of ALLO on PKC activity were seen at any dose tested in mPRε−GFP cells (3nM: 106.34 ± 17.66%, p>0.9999; 10nM: 117.13 ± 19.96%, p=0.9310; 30nM: 88.885 ± 13.12%, p=0.9958; 100nM: 94.011 ± 16.66%, p=0.9983; 300nM: 88.771 ± 16.67%, p>0.9209; Supplemental Figure 5). Thus, in common with ORG, the endogenous NAS, ALLO, activates PKC and PKA signaling dependent upon mPRδ and mPRε, respectively.

### SGE-516 activates mPRδ to increase PKC activity

To further explore the significance of mPRs as effectors for NAS, we tested the effects of the SGE-516, a novel synthetic NAS. SGE-516 has similar efficacy to ALLO as a GABA_A_R PAM and also displays metabotropic effects on GABA_A_R phosphorylation and plasma membrane trafficking.^10,16^ As measured using immunoblotting in mPRδ-GFP, SGE-516 induced significant activation of PKC at 30nM (125.38 ± 11.06%, p=0.0012), 100nM (111.04 ± 10.61%, p=0.0242), and 300nM (142.86 ± 7.59%, p<0.0001), but no activation was observed at lower concentrations (3nM: 100.39 ± 6.22%, p=0.5217; 10nM: 89.857 ± 10.85%, p=0.9964; Figure 6A-B). Consistent with our results from ORG and ALLO treatment, SGE-516 did not modify PKA activity in mPRδ-GFP cells under the same conditions (3nM: 96.916 ± 3.98%, p=0.9985; 10nM: 98.971 ± 6.61%, p>0.9999; 30nM: 93.099 ± 2.69%, p=0.9953; 100nM: 97.858 ± 2.02%, p=0.9228; 300nM: 98.665 ± 4.59%, p=0.8663; Supplemental Figure 6).

**Figure 6.**
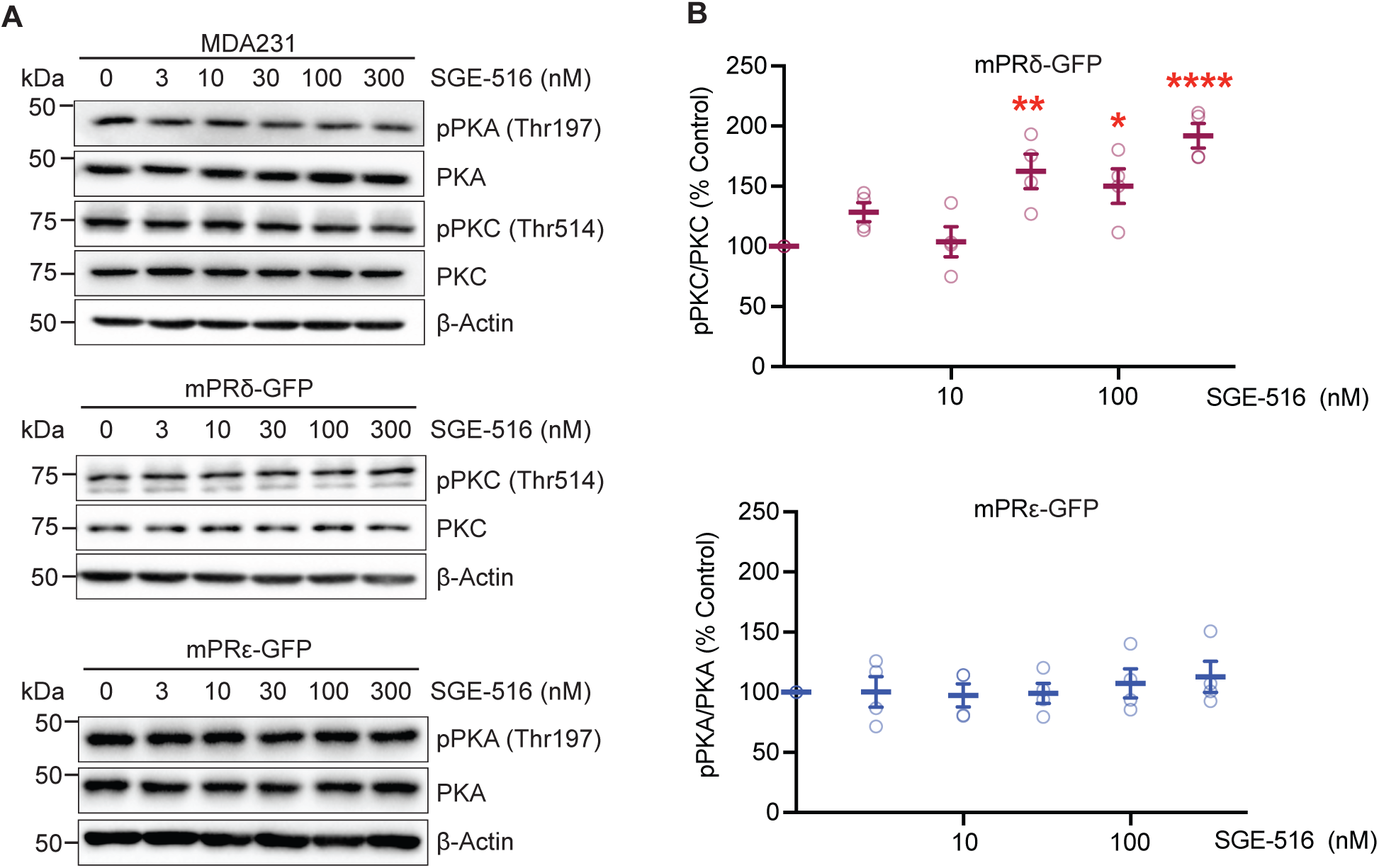
Measuring the effects of SGE516 on mPR signaling. Only cells expressing mPRδ demonstrate PKC activation in response to SGE-516 treatment. **A**. Representative immunoblots for MDA231, mPRδ−GFP, and mPRε-GFP cells treated for 20 minutes with either SGE-516 at 3, 10, 30, 100, and 300nM. Cells were lysed, the proteins resolved on SDS-PAGE and subject to immunblotting**. B**. Densitometry was carried out to quantify the difference between host and mPR expressing cells and between SGE-516 treatment concentrations. The ratio of phosphorylated kinase to total kinase was determined and normalized to control treatment (100%) for each cell type. mPR-GFP cell responses were then normalized to MDA231 means for each SGE-516 concentration. Two-Way ANOVA was used to compare treatment responses in mPRδ−GFP or mPRε−GFP to MDA231 cells. Post hoc comparisons of each SGE-516 concentration were calculated using the Šidák multiple comparison test. * p<0.05, ** p<0.01, *** p<0.001, n=4. In all panels, data represents mean ± SEM.

Similar experiments were then performed in the mPRε-GFP cells. In contrast to our results with ORG (Figure 3) and ALLO (Figure 5), SGE-516 treatment did not increase PKA activation (3nM: 92.279 ± 11.70%, p>0.9999; 10nM: 95.480 ± 9.52%, p>0.9999; 30nM: 98.210 ± 8.41%, p>0.9999; 100nM: 93.685 ± 10.55%, p=0.9981; 300nM: 97.526 ± 11.23%, p=0.9690) or PKC activation (3nM: 79.551 ± 6.53%, p=0.9949; 10nM: 81.896 ± 6.02%, p=0.5457; 30nM: 82.455 ± 7.35%, p>0.9999; 100nM: 88.923 ± 8.26%, p=0.8816; 300nM: 96.873 ± 7.58%, p=0.3577; Figure 6A-B, Supplemental Figure 6) at any dose in the mPRε−GFP. Thus, in contrast to ORG and ALLO, SGE-516 activates mPRδ to increase PKC activity, but not mPRε.

### NAS increase PI3K activity and cAMP accumulation dependent upon mPRδ and mPRε

Our results suggests that mPRδ couples to G_q_ while mPRε couples to G_s_. To directly test this hypothesis, these mPR cell lines were transfected with luminescent reporters to indirectly measure PI3K, the principal effector of G_q_, or elevations in cAMP accumulation, which reflect activation of G_s_. 72 hours following transfection, cells were treated with increasing concentrations of ORG, ALLO, or SGE-516. Luminescence was then measured and compared between control and various compound concentrations. In mPRδ−GFP cells, ORG, ALLO, and SGE-516 significantly increased G_q_-coupled luminescence with similar EC_50_ values of 1.7nM, 11.3nM and 3.7nM, respectively (Figure 7A,C). For mPRε-GFP cells, significant elevations in G_s_-coupled luminescence were seen with ORG and ALLO with EC_50_s of 11.3nM and 2.5nM, respectively (Figure 7B-C). In contrast, SGE-516 did not display a concentration-dependent response in mPRε−GFP. Collectively our results suggest that mPRδ couples to G_q_, whilst mPRε signals via G_s_. In addition, these assays provided further evidence that SGE-516 is a potent agonist for mPRδ but not mPRε.

**Figure 7.**
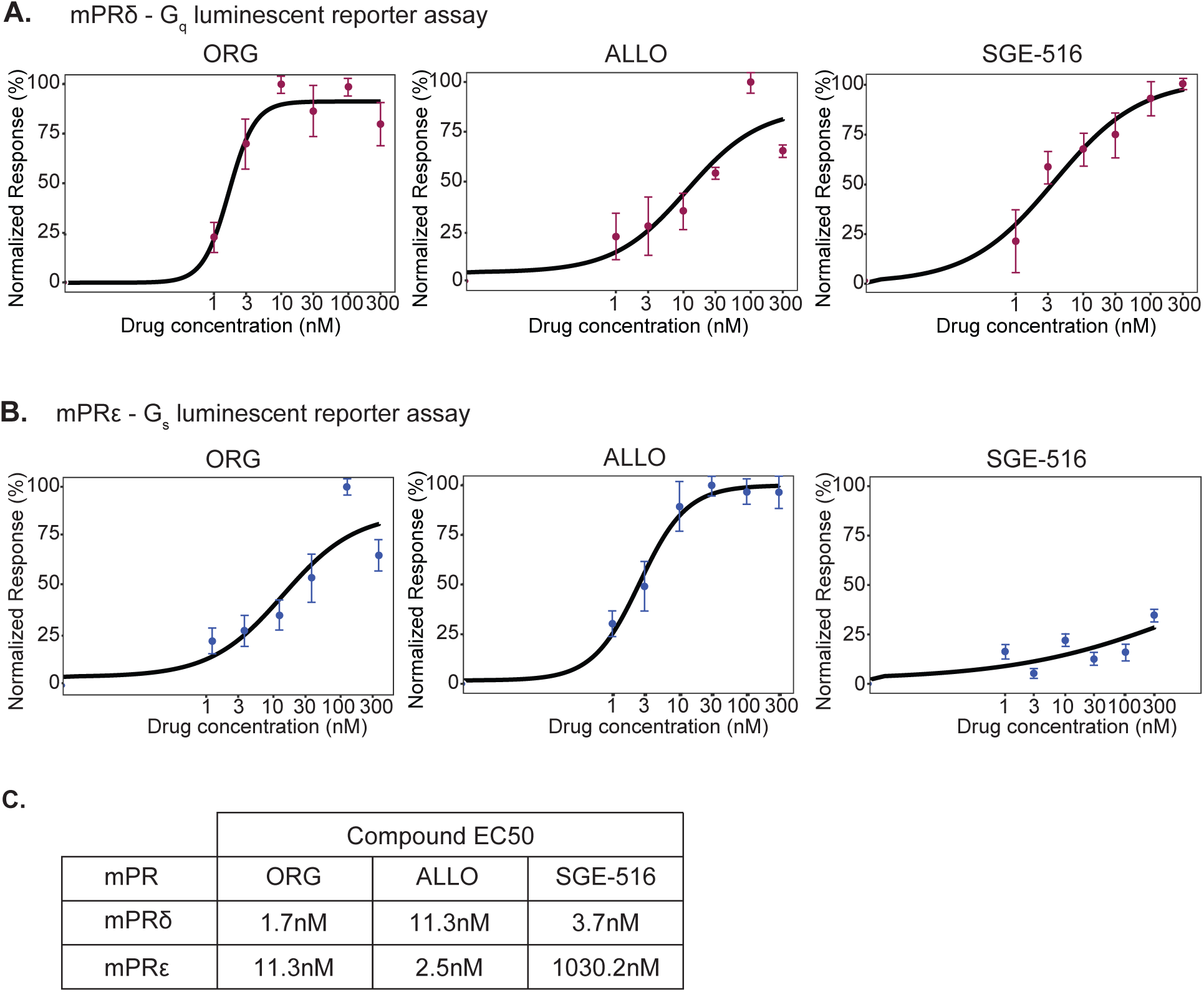
Analyzing the effects of NAS on G_q_ and G_s_ activation. **A.** Dose response curves for ORG, ALLO and SGE-516, using a G_q_ reporter assay in mPRδ−GFP cells. Cells were exposed to 1-300nM ORG, ALLO or SGE-516 for 6 hours. Data were normalized to the minimum and maximum luminescent values curves fitted to calculate EC_50_ values (*n*=4). **B**. Dose response curves for ORG, ALLO and SGE-516, using a G_s_ reporter assay in mPRε-GFP. Cells were exposed to 1-300nM ORG, ALLO or SGE-516 for 6 hours. Data were normalized to the minimum and maximum luminescent values curves fitted to calculate EC_50_ values (*n*=4). **C.** EC_50_ values were determined for ORG, ALLO and SGE-516 in both mPRδ−GFP and mPRε-GFP cells.

### SGE-516 and ORG increase PKC activity in the hippocampus of female but not male C57BL/6 mice

To test the significance of our studies in cell lines for signaling in the brain, we examined the expression levels of mPRδ and mPRε in the forebrains of 8–12-week-old C57BL/6 wild-type (WT) male and female mice. In the absence of suitable antibodies, we measured the levels of the respective mRNAs using real-time quantitative polymerase chain reaction (RT-qPCR) with subtype specific primers. The relative levels of mPR mRNAs were then compared by reference to GAPDH and the results were then expressed as arbitrary units, as described previously.^16^ In female mice the levels of mPRδ mRNA were 815.0 ± 8.5% compared to males (100.0 ± 4.7%, p<0.0001; Figure 8A). Likewise, 68% higher levels of mPRε were found in females compared to males (Females: 168.0% ± 2.3%; Males: 100 ± 0.3%, p<0.0001; Figure 8A).

**Figure 8.**
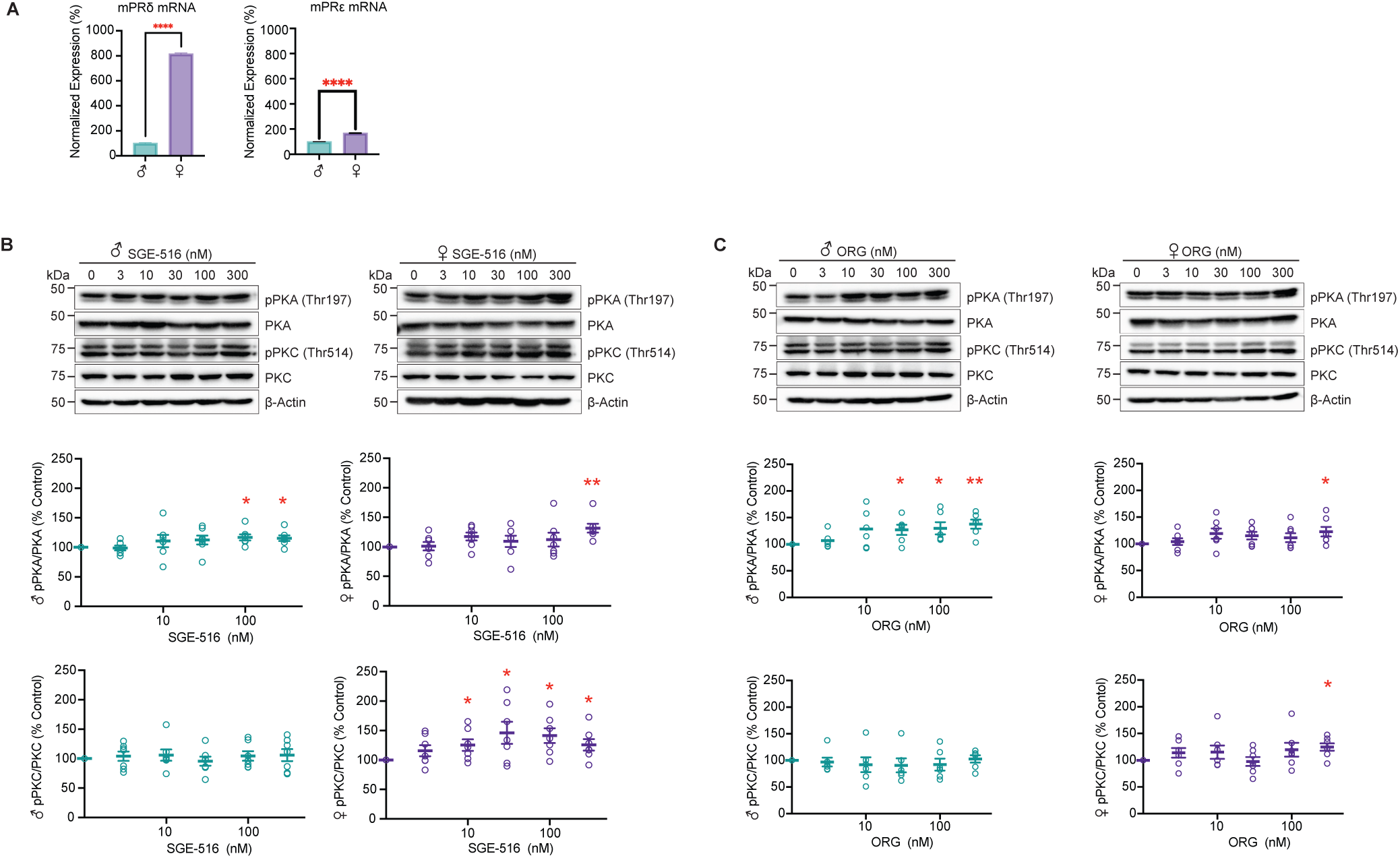
Examining the effects of SGE-516 and ORG on PKC activity in hippocampal slices. **A.** RT-qPCR was performed with forebrain from adult WT male or female mice. Expression was normalized to males (100%) (*n*=4 groups of 7 mice per group). Error bars represent the SEM. Welch’s t-test was used to compare males to females. **B**. Immunoblots for hippocampal slices from adult WT male or female mice treated for 20 minutes with SGE-516 at 3, 10, 30, 100, and 300nM. Hippocampal slices were lysed, the proteins resolved on SDS-PAGE and subject to immunblotting. Densitometry was carried out to quantify the difference between control and SGE-516 concentrations. The ratio of phosphorylated kinase to total kinase was determined and normalized to control treatment (100%). **C**. Immunoblots for hippocampal slices from adult WT male or female mice treated for 20 min with ORG at 3, 10, 30, 100, and 300nM. Hippocampal slices were lysed, the proteins resolved on SDS-PAGE, and subject to immunblotting. Densitometry was carried out to quantify the difference between control and ORG concentrations. The ratio of phosphorylated kinase to total kinase was determined and normalized to control treatment (100%). Lines and error bars represent mean ± SEM (*n*=7). Welch’s t-test was used to compare control treatment to each SGE-516 or ORG concentration. * p<0.05, ** p<0.01, **** p<0.0001.

To examine the physiological significance of our RT-qPCR results, we prepared hippocampal brain slices from 8–12-week-old male and female C57BL/6 mice. Following at least 60 min of recovery, slices were treated with various concentrations of SGE-516 for 20 min. Then immunoblots were used to assess changes in PKC and PKA phosphorylation as a measure of kinase activation. Significantly increased PKA activation was observed in both male and female WT mice following treatment with SGE-516. Males showed increased PKA activity after treatment with 100nM and 300nM SGE-516 compared to the effect seen in females, where increased activation was only observed following 300nM SGE-516 (Males: 100nM: 116.82 ± 5.27% p=0.032, 300nM: 115.53 ± 4.78%, p=0.0418; Females: 300nM: 131.49 ± 7.54%, p=0.0058; Figure 8B-C). No changes in PKA activation were observed at any other SGE-516 concentrations (Males: 3nM: 98.862 ± 4.08%, p=0.7895, 10nM: 110.67 ± 10.47%, p=0.3475, 30nM 112.43 ± 7.62%, p= 0.1541; Females: 3nM: 100.86 ± 7.14%, p=0.9079, 10nM: 117.34 ± 6.66%, p=0.0504, 30nM 109.23 ± 9.55%, p=0.3712, 100nM: 112.29 ± 11.72%, p=0.3347).

In contrast to PKA activation, significant activation of PKC following SGE-516 treatment was only observed in female WT mice. This was observed following doses as low as 10nM of SGE-516 (3nM: 115.66 ± 9.52%, p=0.1509; 10nM: 125.26 ± 9.83%, p=0.0423; 30nM: 146.23 ± 18.82%, p=0.0494; 100nM: 141.35 ± 12.59%, p=0.0167; 300nM: 125.72 ± 9.83%, p=0.0398; Figure 8B,D). No changes in PKC activation were observed in male WT mice (3nM: 104.05 ± 8.02%, p=0.6315; 10nM: 105.92 ± 9.62%, p=0.5608, 30nM: 95.654 ± 7.79%, p=0.5971; 100nM: 104.44 ± 8.08%, p=0.6026; 300nM: 106.00 ± 10.29%, p=0.5812).

In addition to NAS, we also assessed the effects that the mPR agonist ORG has on intracellular signaling in acute brain slices. Importantly, in contrast to ALLO or SGE-516, ORG does not exert any efficacy at GABAARs.^16^ Like SGE-516, ORG treatment led to a significant increased PKA activation in both male and female WT mice. Males showed increased PKA activity after treatment with 30, 100nM and 300nM ORG. In females, increased PKA activation was only observed following 300nM ORG (Males: 30nM: 127.02 ± 9.58% p=0.037, 100nM: 129.84 ± 11.48% p=0.048, 300nM: 137.54 ± 8.66%, p=0.0075; Females: 300nM: 122.80 ± 9.04%, p=0.0452; Figure 8E-F). No changes in PKA activation were observed at any other ORG concentrations (Males: 3nM: 106.71 ± 5.77%, p=0.2974, 10nM: 128.36 ± 14.38%, p=0.1056; Females: 3nM: 104.23 ± 6.20%, p=0.5208, 10nM: 119.19 ± 9.62%, p=0.0933, 30nM 115.33 ± 7.05%, p=0.0726, 100nM: 111.49 ± 8.49%, p=0.2247).

In contrast to PKA activation, significant activation of PKC was only observed in female WT mice following treatment with 300nM ORG (124.76 ± 6.94%, p=0.0118; Figure 8E,G). No changes in PKC activation were observed at lower doses of ORG in female WT mice (3nM: 113.91 ± 9.06%, p=0.1757, 10nM: 115.05 ± 12.60%, p=0.2774; 30nM: 97.70 ± 8.39%, p=0.7936; 100nM: 119.65 ± 12.86%, p=0.1773) or in male WT mice (3nM: 97.05 ± 8.44%, p=0.7410; 10nM: 91.91 ± 14.05%, p=0.5899, 30nM: 90.64 ± 13.13%, p=0.5216; 100nM: 92.22 ± 11.29%, p=0.7022; 300nM: 102.83 ± 6.99%, p=0.7022).

Collectively these studies demonstrate that SGE-516 selectively activates PKC in females via activation of mPRδ, which correlates with the 8-fold elevation in mPRδ mRNA expression in female mice.

## Discussion

NAS that act as GABA_A_R PAMs are potent endogenous modulators of neuronal excitability and this activity underlies their effects on behavior and their therapeutic efficacy in PPD and CDKL5 deficiency disorder.^4,5,20,41–44^ In this study we explored the role that mPRs, a family of metabotropic receptors that bind P4 and other progestins, play as effectors for NAS, specifically focusing on the mPRδ and ε isoforms, which are expressed in the nervous system and peripheral tissues.

To do so, we created MDA231 cell lines that stably express mPR isoforms modified at the C-terminus with GFP. We first explored the subcellular localization of mPRδ and mPRε in each cell line, which revealed high levels of accumulation within the ER. However, a proportion of each mPR was able to accumulate in other compartments within the secretory or endocytic pathways, or the plasma membrane. In contrast to this, mPRα, β and γ have been reported to be retained within the ER when expressed in some cells.^19,29^ The discrepancies may reflect subtype specific differences in the subcellular localization of individual mPR isoforms or methodological variations between studies.

Live TIRF imaging was also used to examine mPR expression and stability on the plasma membrane. Consistent with our data using immunostaining, robust fluorescence was seen for both mPRs using TIRF suggesting accumulation within 100nM of the plasma membrane. Rapid internalization of mPRδ but not mPRε was seen with pan-mPR agonist ORG, a classical property of canonical GPCRs which regulates the duration of signaling and effector specificity.^40^ Classical studies have revealed that agonist activation of GPCRs promotes recruitment of β-arrestins and subsequent clathrin dependent endocytosis. The respective GPCR/β-arrestin complexes signal within the endocytic pathway, often via distinct effectors to those activated on the plasma membrane.^45^ Clearly, while further studies are required to assess the significance of mPRδ internalization, it may act to regulate the duration and diversity of NAS signaling.

We then sought to assess if ORG increased the activity of known protein kinase effectors for G-proteins in our cell lines focusing on PKC, PKA and Src that reflect activation of G_q_, G_s_ and G_i_ respectively. ORG potently activated PKC via mPRδ and increased PKA activity via mPRε with effects being evident at <30nM. These findings were replicated following an acute treatment of ALLO. However, SGE-516 treatment increased PKC but not PKA activity, suggesting SGE-516 activates mPRδ but not mPRε.

To further characterize mPR signaling we employed G_q_ and G_s_ reporter assays. The results from these reporter assays confirmed that mPRδ behaves as a G_q_-coupled GPCR to activate PKC while mPRε behaves as a G_s_-coupled GPCR to activate PKA and that SGE-516 activation of mPRs was limited to mPRδ. We were also able to calculate the EC_50_ of each NAS using our luminescent reporter assays. These ranged from 1.7 to 11.3nM for ORG and ALLO for both mPRδ and mPRε. In contrast, SGE-516 has an EC_50_ of 3.7nM for mPRδ, but >1μM for mPRε. The EC_50_ of these compounds for mPRδ and mPRε are within the endogenous concentrations of ALLO that have been reported.^2,46^ This suggests that the brain is sensitive to low concentrations of NAS and these may activate mPR signaling pathways.

This is the first evidence that mPRδ and mPRε display divergent signaling via PKC and PKA. Previous work has demonstrated that activation of ERK, AKT, and MAPK occur downstream of mPRs.^47,48^ For example, in MDA231 cells that were transiently transfected to express mPRδ, a significant increase in the phosphorylation of ERK was observed following progesterone and ALLO treatment.^17^ ERK, MAPK, and AKT are downstream kinases of PKC and PKA, therefore our work is consistent with these previous findings. Additionally, we confirmed involvement of G_q_ and G_s_ signaling both by immunoblot and luminescent reporter assays, thereby reinforcing our findings. Importantly, both PKC and PKA are known to be critical for the effects of some NAS on GABA_A_R phosphorylation and trafficking. In acute brain slices, the activity of PKC and PKA is necessary for the increase in phosphorylation of the GABA_A_R β3 subunit and for the increase in PM expression of the α4 and β3 subunits seen following treatment with either NAS, like ALLO or SGE-516, or the mPR agonist, ORG.^10,15,16,49^ This suggests that signaling through mPRs could act as additional mechanisms mediating the effects of NAS on neuronal signaling, which may contribute to their therapeutic efficacy as anti-depressant and anticonvulsant therapies.

To assess the significance of our studies in cell lines, we examined the expression levels of mPRδ and mPRε in the mouse brain revealing that both mPRδ and mPRε are expressed at higher levels in females compared to males. Treatment of hippocampal slices from male and female WT C57Bl/6 mice with SGE-516 led to increased PKA activation in both sexes. This contrasts with our results in the MDA231 cell lines where we saw changes in PKC activation in only mPRδ-GFP cells. This discrepancy may reflect the complexity of cell types between experimental preparations, and the presence of mPRα, mPRβ, and mPRγ, as the encoding mRNAs are expressed in the hippocampus.^17^ SGE-516 may act on one or more of these mPRs to modulate cAMP accumulation. Additionally, it is possible that these effects are mediated by other as yet unknown receptors for NAS.

Following treatment with SGE-516, we also demonstrated robust and potent activation of PKC in brain slices from female, but not male mice. Similar activation of PKC in only female WT mice following treatment with the pan-mPR agonist ORG suggests that the effects of SGE-516 on PKC activation is due to signaling via mPRs as opposed to its effects as a GABA_A_R PAM. Additionally, the preferential activation of PKC only in female WT mice correlates with our results demonstrating higher expression of mPRδ in the female brain. Finally, our results also suggest that female WT mice may require a higher concentration of ORG to induce PKA activation compared to male WT mice highlighting other potential sex-specific effects in mPR signaling in the brain.

Collectively our work demonstrates that NAS like ALLO and SGE-516 exert potent effects on G_q_ signaling via mPRδ, and these effects may be sex-specific. In addition, ALLO also exerts potent effects on G_s_ via mPRε. Thus, it is possible that mPRs may mediate effects of some NAS on neuronal excitability and behavior, highlighting the opportunity for developing specific ligands for these receptors. Additionally, NAS are dysregulated in many neuropsychiatric diseases including postpartum depression, major depressive disorder, anxiety, and schizophrenia.^50^ Therefore, it is essential to gain a better understanding of the expression and mPR-mediated signaling that occurs in the brain in response to NAS to develop new pharmacological agents to treat these and other diseases.

### Limitations of current study

This study demonstrates that NAS like ALLO and SGE-516 lead to sex-specific activation of kinases downstream of mPRδ and mPRε activation. However, there are some limitations to our study that we would like to discuss. First, our studies assess the downstream signaling of mPRδ and mPRε in MDA231cells which are not representative of mPR signaling within the brain. While we validated our findings in acute hippocampal slices to determine the role of mPRδ and mPRε signaling in the brain, but we did not assess how the expression of other mPRs (mPRα, mPRβ, and mPRγ) influences downstream kinase activation. The downstream signaling of mPRα, mPRβ, and mPRγ in isolation as well as in the brain must therefore be further examined to determine a more complete picture of how NAS signal through mPRs to affect neural function. Additionally, while we verify that the murine brain expresses mPRδ and mPRε via RT-qPCR, we were unable to validate protein or cell-type specific expression of these mPRs. This is partially due to a lack of suitable antibodies against these mPRs and should be further explored in the future to better understand mPR signaling in the brain. Finally, the current study focused on the biochemical signaling pathways downstream of NAS activation of mPRδ and mPRε and did not examine electrophysiological or behavioral effects of NAS signaling through mPRs. Therefore, future studies are needed to address how NAS like ALLO and SGE-516 may signal through mPRδ and/or mPRε to contribute to the established anti-convulsant and anti-depressant effects of NAS.

## Acknowledgements

PAD and SJM are supported by National Institutes of Health (NIH) – National Institute of Neurological Disorders and Stroke grants NS087662 (SJM), NS081986 (SJM), NS108378 (PAD and SJM), NS101888 (SJM), NS103865 (SJM), and NS111338 (SJM) and NIH – National Institute of Mental Health grant MH118263 (SJM) and MH097446 (PAD and SM. Manasa Parakala and Jon Madison from Sage Therapeutics for providing reagents, expertise, funding, and manuscript editing.

## Author Contributions

AHSL, BM, JSD, SS and JLS performed research. AHSL, JLS, PD, and SJM designed experiments and wrote the manuscript.

## STAR ★ METHODS

## KEY RESOURCES TABLE

| REAGENT or RESOURCE | SOURCE | IDENTIFIER |
| --- | --- | --- |
| **Antibodies** |  |  |
| Mouse β-actin | Sigma | A1978 |
| Rabbit PKA | Cell Signaling | 5842S |
| Rabbit pPKA | Cell Signaling | 5661S |
| Rabbit PKC | Cell Signaling | 2056S |
| Rabbit pPKC | Cell Signaling | 9379S |
| Rabbit Src | Cell Signaling | 2123S |
| Rabbit pSrc | Cell Signaling | 6943S |
| Chicken GFP | Abcam | ab13970 |
| Mouse BAP31 | Abcam | ab112993 |
| Goat anti-chicken Alexa Fluor 488 | ThermoFisher | A-11039 |
| Goat anti-mouse Alexa Fluor 568 | ThermoFisher | A-11004 |
| **Chemicals, peptides, and recombinant proteins** |  |  |
| Allopregnanolone | Sigma | P8887 |
| Org OD 02-0 | Axon MedChem | 2085 |
| SGE-516 | Sage Therapeutics | N/A |
| **Critical commercial assays** |  |  |
| Luciferase Assay Reagent | Promega | E1483 |
| Luciferase Cell Culture Lysis Reagent | Promega | E1500 |
| **Experimental models: cell lines** |  |  |
| MDA-MB-231 HTB-26 cells | ATCC | HTB-26 |
| **Experimental models: Organisms/strains** |  |  |
| Mice: C57BL/6J strain | JAX | Strain #000664 |
| **Recombinant DNA** |  |  |
| FHRE-Luc plasmid | Addgene | 1789 |
| pGL4_CRE-CMVmin-luc2 plasmid | Addgene | 194384 |
| **Software and algorithms** |  |  |
| ImageJ/Fiji | NIH | https://imagej.nih.gov/ij/ |
| GraphPad Prism 10 | GraphPad Software | https://imagej.nih.gov/ij/ |
| Softmax Pro Software for Flexstation 3 Multi-Mode Microplate Reader | Molecular Devices | Flex3 |

## RESOURCE AVILABILITY

### Lead contact

Further information and requests for resources should be directed to and will be fulfilled by the lead contact, Professor Stephen Moss

### Materials availability

This study did not generate new unique reagents.

### Data and code availability

- Original immunoblotting and microscopy data reported in the paper will be shared by the lead contact upon request.
- No original code was developed by this study.
- Any additional information required to reanalyze the data reported in this paper is available from the lead contact upon request.

## EXPERIMENTAL MODELS AND SUBJECT DETAILS

### Animals

Animal housing protocols and surgical procedures were performed according to protocols approved by the Institutional Animal Care and Use Committee of Tufts Medical Center (IACUC). C57/BL6 Mice (4-12wk) were maintained at 22°C2 on a standard 12-h light/dark cycle with free access to food and water.

### Cell lines

MDA-MB-231 HTB-26 cells were purchased from ATCC (https://www.atcc.org/products/htb-26).

## METHOD DETAILS

### Animals

Animal studies were performed according to protocols approved by the Institutional Animal Care and Use Committee (IACUC) of Tufts Medical Center. 8–12-week-old C57BL/6 male and female mice were kept on a 12-hour light/dark cycle with *ad libitum* access to food and water. Mice were sacrificed by being anesthetized with isoflurane and then rapidly decapitated.

### Tissue Culture

Tissue culture was carried out as previously described.^51^ Briefly, MDA-MB-231 cells were obtained from ATCC (Manassas, VA) and maintained in DMEM supplemented with 10% charcoal stripped fetal bovine serum (FBS) and penicillin/streptomycin. All cells were passaged at 90% confluency, for maintenance. The cells were used for experiments at a passage number no higher than 20, and for stable cell line generation, the lowest possible passage number was used.

### Cell Line Production

MDA-MB-231 cells were transduced with lentivirus for mouse mPRδ and mouse mPRε with a c-terminal GFP tag (Origene, mPRδ: MR216535L4V, mPRε: MR215906L4V). The cells were exposed to 10^5^ genomic copies per cell for 24 hours in a 24-well plate, followed by a media change to remove the lentivirus. The transduced cell lines were expanded over two passages into 10cm dishes. At 90% confluency, the cells were trypsinized, washed and resuspended in PBS. 5000 individual GFP-positive events (cells) were selected by FACS and plated in a 24-well plate well. This process was repeated twice. During the third FACS sort, individual GFP-positive cells were sorted into each well of a 96-well plate. These cells were allowed to grow for several weeks with daily visual monitoring of growth-rate and GFP-expression. The clone that demonstrated the highest GFP expression and favorable growth rate was selected for expansion (Supplemental Figure 1).

### Acute Cortical/Hippocampal Slices

Acute slices were prepared as previously described.^52^ Briefly, 310μm thick coronal forebrain slices were obtained from wild type mice. The slices were recovered in oxygenated aCSF for 60-90 min. The slices were then transferred to a custom microdialysis chamber in oxygenated aCSF containing varying concentrations of SGE-516 for 20 min. The slices were then removed from the chamber and snap frozen for immunoblotting.

### Western Blotting

Western blots were carried out as previously described.^53^ Briefly, for cultured cells, the cell monolayer was washed with PBS and then the cells were lysed in RIPA buffer containing protease and phosphatase inhibitors. Acute brain slices were rapidly thawed and lysed in RIPA buffer containing protease and phosphatase inhibitors. Proteins were quantified by Bradford assay. Samples were diluted to the same concentration in RIPA buffer and 2X sample buffer added. Samples were boiled for 5 min at 95 °C before being loaded onto polyacrylamide gels for sodium dodecyl sulfate–poly acrylamide gel electrophoresis (SDS-PAGE). Proteins were then transferred onto nitrocellulose membranes, blocked in 5% milk for 1 hour, and probed with primary antibodies overnight. The membranes were washed and probed with appropriate HRP-conjugated secondary antibodies and developed with enhanced chemiluminescent (ECL) substrate in a Bio-Rad ChemiDoc Imager. Where possible all replicates were run on the same gel.

### Biochemical Assays

G_q_ and G_s_ reporter assays were performed as previously described.^54–57^ Briefly, clonal stably expressing mPRδ- and mPRε-expressing cells were grown to 90% confluency, harvested, counted using a CASY cell counter. 10μg of G_q_ or G_s_ reporter DNA was transfected into 1x10^6^ cells using Lipofectamine 3000 transfection reagent (G_q_: FHRE-Luc, G_s_: pGL4_CRE-CMVmin-luc2). The FHRE-Luc plasmid used to assess G_q_ activation was a gift from Michael Greenberg (Addgene plasmid # 1789 ; http://n2t.net/addgene:1789; RRID:Addgene_1789). The pGL4_CRE-CMVmin-luc2 plasmid used to assess G_s_ activation was a gift from Michael Wehr (Addgene plasmid # 194384 ; http://n2t.net/addgene:194384; RRID:Addgene_194384). The transfected cells were plated in a 96-well plate at 1x10^4^ cells per well. The cells were grown for 72 hours, and then treated with varying concentrations of NAS for 6 hours, or positive control compounds (10μM PDBu or 10μM forskolin for G_q_ or G_s_ assays respectively). The cells were lysed in 20μl per well of Luciferase Cell Culture Lysis Reagent (Promega) for 20 min. 80μl of Bright-Glo Luciferase Assay reagent (Promega) was added to each well and incubated for 2 min. The luminescence of each well was measured using a Flexstation plate reader (Molecular Devices). Only data where the mean positive control well values were at least double the mean luminescence of the untreated controls were used. Minimum and maximum luminescence values for each compound were scaled between 0 and 100 respectively and used to plot dose-response curves.

### Immunocytochemistry

Immunocytochemistry was carried out as previously described.^58^ Briefly, cultured MDA231/mPR-GFP cells were washed with PBS, fixed in 4% paraformaldehyde (PFA, Electron Microscopy Services) in PBS, and permeabilized in block solution consisting of 1X PBS with 0.5% Triton X-100, 5% (w/v) Bovine Serum Albumin (BSA). Primary and secondary antibodies (as described above) were prepared in block solution and incubated with the cells for 1 hour at room temperature in the dark. The cells were washed in 1X PBS and mounted on glass slides using Fluoromount-G. The cells we imaged using a Nikon Eclipse Ti (Nikon Instruments, Melville, NY, United States) confocal microscope using a 60x oil immersion objective lens.

For analyzing colocalization, the BIOP JACoP plugin for ImageJ/Fiji was used. Images of mPR-GFP cells were auto-thresholded using the Otsu Thresholding method. The area of GFP signal and BAP31 signal were then quantified individually and the overlapping area between the two fluorophores was also calculated. This number was reported as the percent of GFP signal colocalizing with or without BAP31 signal.

### TIRF Microscopy

TIRF microscopy was carried out as previously described.^15^ Briefly, MDA231 cells were plated in 35mm glass-bottom dishes and grown to approximately 50% confluency. The dishes were removed from the incubator and the media was replaced with HEPES-buffered aCSF. The cells were placed in an imaging chamber heated to 37°C attached to a Nikon Eclipse Ti Inverted TIRF Microscope (Nikon Instruments). The GFP signal was imaged for 5 min to establish a baseline using a 60x oil-immersion objective. The cells were then perfused with aCSF containing 100nM ORG and imaged for a further 20 min. The total fluorescence was normalized to the mean baseline value.

### QUANTIFICATION AND STATISTICAL ANALYSIS

All results are expressed as mean ± the standard error the mean (SEM). For the MDA231 cell experiments, all compound response data was first normalized to control treatment (100%) within each cell line and then normalized to the response in the host MDA231 cells. A two-way ANOVA was then used to compare kinase activation responses between the transduced mPR-GFP cells and the host MDA231 cells. Post-hoc comparisons of each compound concentration were calculated using the Šidák multiple comparison test. For acute hippocampal slice experiments, Welch’s t-test was used to compare each concentration of SGE-516 or ORG to the control treatment. All replicates are independent biological replicates. For all MDA231 cell experiments, *n* = 4 was used from separate cell passages. For acute hippocampal slice experiments, *n* = 7 was used.

