## Supplemental Data Figures for "Neuroactive steroids activate membrane progesterone receptors to induce sex specific effects on protein kinase activity"

**Supplemental Figure 1. FACs sorting of viral infected MDA231 cells. A**. Representative plots of Fluorescence Activated Cell Sorting (FACS) based enrichment of GFP-positive MDA231 cells transduced with mPRδ-GFP or mPRε-GFP. 5000 GFP-positive cells were selected by forward and side scatter (to select for individual cells), followed by GFP (FITC) intensity, and placed in a 24-well plate well. **B.** Representative plots of FACS based single-cell selection of GFP-positive MDA231 cells transduced with mPRδ-GFP or mPRε-GFP. Single GFP-positive cells were selected by forward and side scatter (to select for individual cells), followed by two GFP (FITC) intensity sorts. A single GFP-positive cell was placed in each well of a 96-well plate for expansion.

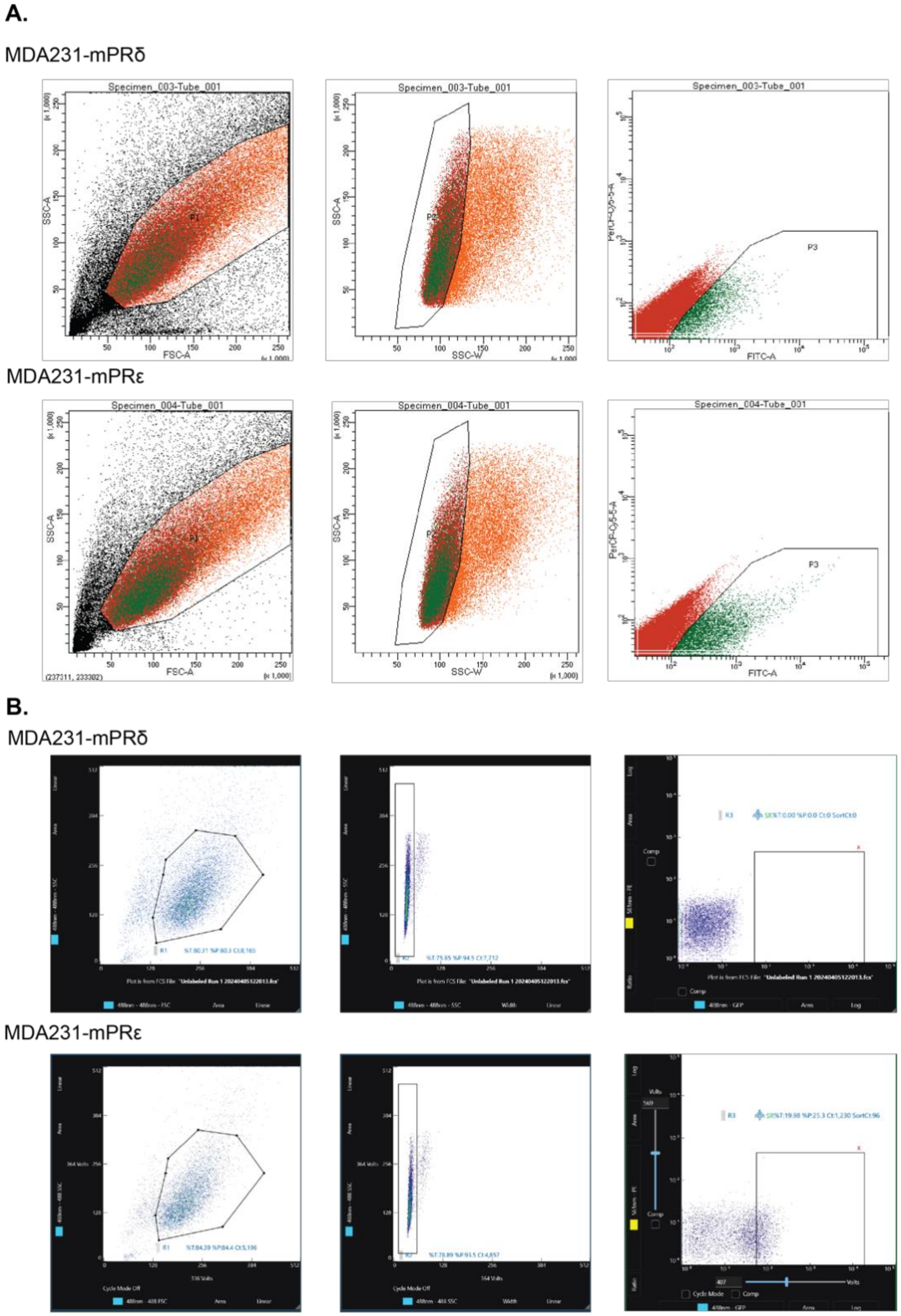

**Supplemental Figure 2. TIRF imaging of mPRδ-GFP and mPRε-GFP cells.** Cultures were fixed, permeabilizied and immunostained with actin together with GFP antibodies and subject to TIRF imaging. Scale bar represents 20μm.

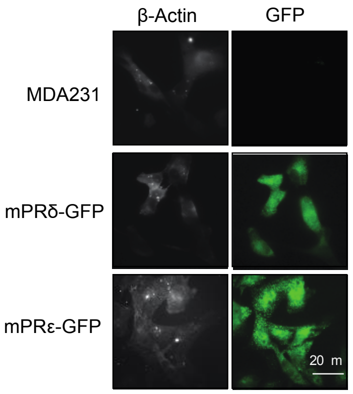


**Supplemental Figure 3. Examining the effects of ORG on PKC activity in cell lines expressing mPRs. A**. Representative immunoblots for MDA231-mPRδ cells treated with ORG (300nM) for 5, 10, 15, or 20 minutes. Cells were lysed, the proteins resolved on SDS-PAGE and subject to immunoblotting Densitometry was carried out to quantify the difference in PKC activation following 300nM ORG treatment after either 5, 10, 15, or 20 minutes. The ratio of phosphorylated kinase to total kinase was determined and normalized to PKC activation at baseline (100%) without ORG treatment (*n*=4).

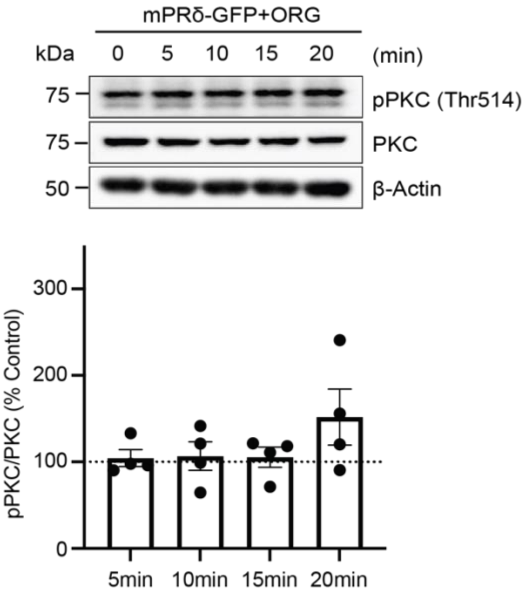

**Supplemental Figure 4: Examining the effects of ORG on protein kinase activity in cell lines. A**. Representative immunoblots for MDA231, MDA231-mPRδ, and MDA231-mPRε cells treated for 20 minutes with increasing concentrations of ORG. Cells were lysed, the proteins resolved on SDS-PAGE, and subject to immunoblotting **B**. Densitometry was carried out to quantify the difference between ORG treatment concentrations and control treatment in MDA231 cells. The ratio of phosphorylated kinase to total kinase was determined and normalized to control treatment (100%) for each cell type (*n*=4). **C**. Densitometry was carried out to quantify the difference between host and mPRδ expressing cells and between ORG treatment concentrations. The ratio of phosphorylated kinase to total kinase was determined and normalized to control treatment (100%) for each cell type (*n*=4). **D**. Densitometry was carried out to quantify the difference between host and mPRε expressing cells and between ORG treatment concentrations. The ratio of phosphorylated kinase to total kinase was determined and normalized to control treatment (100%) for each cell type (*n*=4). Two-Way ANOVA was used to compare treatment responses in MDA231-mPRδ or MDA231-mPRε to MDA231 cells. Post hoc comparisons of each ORG concentration were calculated using the Šidák multiple comparison test.

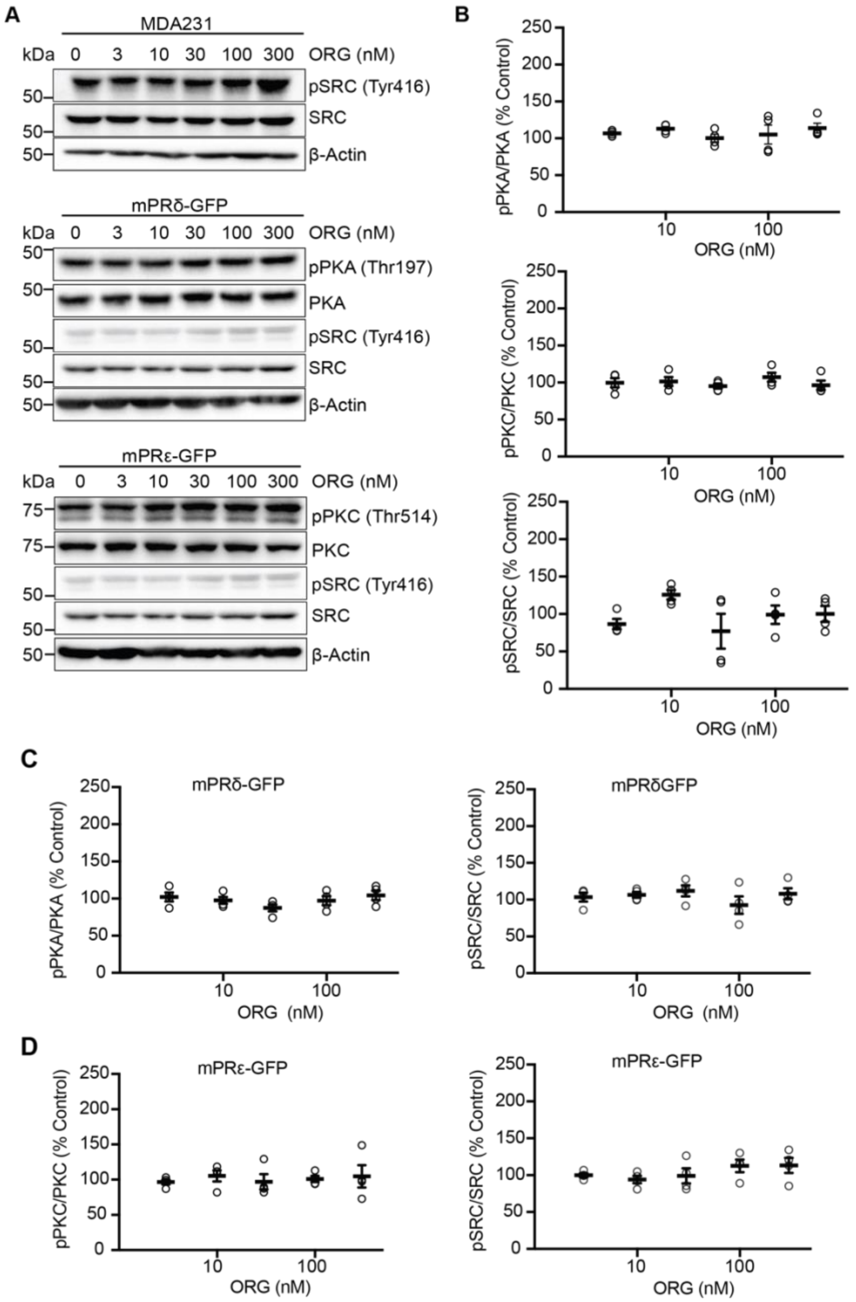

**Supplemental Figure 5. Examining the effects of ORG on protein kinase activity in cell lines. A**. Representative immunoblots for MDA231, MDA231-mPRδ, and MDA231-mPRε cells treated for 20 minutes with ALLO. Cells were lysed, the proteins resolved on SDS-PAGE, and subject to immunoblotting. **B.** Densitometry was carried out to quantify the difference between host and mPR expressing cells and between ALLO treatment concentrations. The ratio of phosphorylated kinase to total kinase was determined and normalized to control treatment (100%) for each cell type (*n*=4). Two-Way ANOVA was used to compare treatment responses in MDA231-mPRδ or MDA231-mPRε to MDA231 cells. Post hoc comparisons of each ALLO concentration were calculated using the Šidák multiple comparison test.

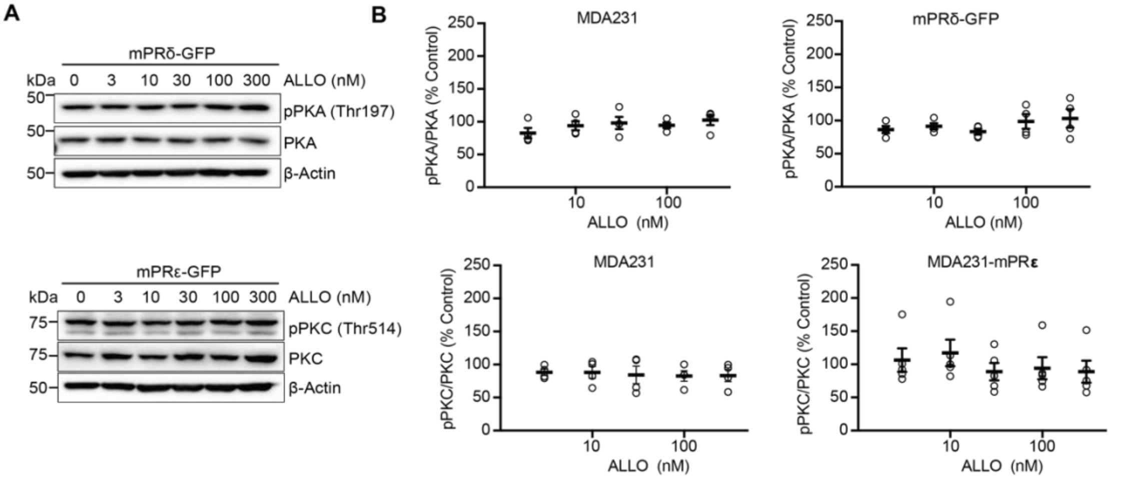


**Supplemental Figure**

**Supplementary Figure 6. Examining the effects of SGE-516 on protein kinase activity in cell lines. A**. Representative immunoblots for MDA231, MDA231-mPRδ, and MDA231-mPRε cells treated for 20 minutes with SGE-516 at 3, 10, 30, 100, and 300nM. Cells were lysed, the proteins resolved on SDS-PAGE, and subject to immunoblotting. **B.** Densitometry was carried out to quantify the difference between host and mPR expressing cells and between SGE-516 treatment concentrations. The ratio of phosphorylated kinase to total kinase was determined and normalized to control treatment (100%) for each cell type (*n*=4). Two-Way ANOVA was used to compare treatment responses in MDA231-mPRδ or MDA231-mPRε to MDA231 cells. Post hoc comparisons of each SGE-516 concentration were calculated using the Šidák multiple comparison test.

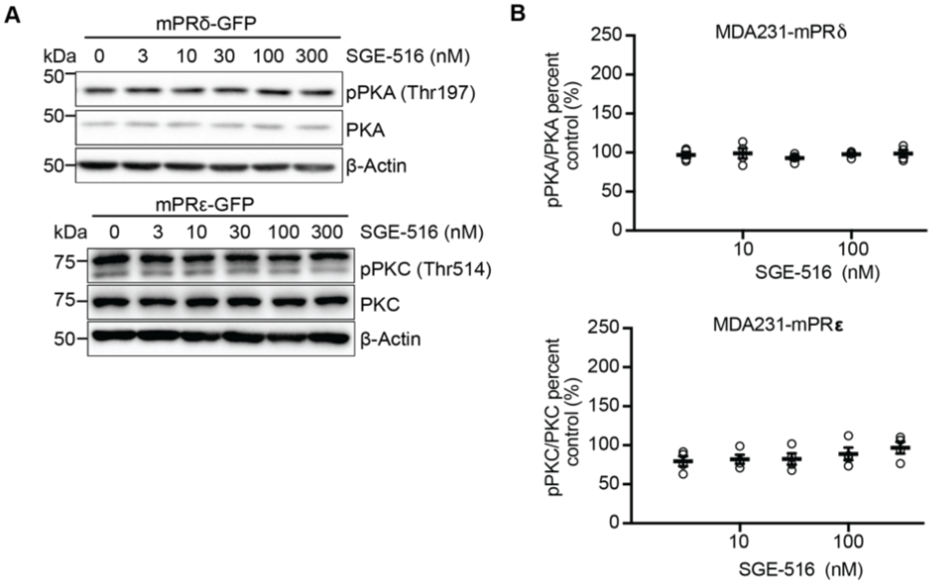


=

**Supplemental Figure 1. FACs sorting of viral infected MDA231 cells. A**. Representative plots of Fluorescence Activated Cell Sorting (FACS) based enrichment of GFP-positive MDA231 cells transduced with mPRδ-GFP or mPRε-GFP. 5000 GFP-positive cells were selected by forward and side scatter (to select for individual cells), followed by GFP (FITC) intensity, and placed in a 24-well plate well. **B.** Representative plots of FACS based single-cell selection of GFP-positive MDA231 cells transduced with mPRδ-GFP or mPRε-GFP. Single GFP-positive cells were selected by forward and side scatter (to select for individual cells), followed by two GFP (FITC) intensity sorts. A single GFP-positive cell was placed in each well of a 96-well plate for expansion.

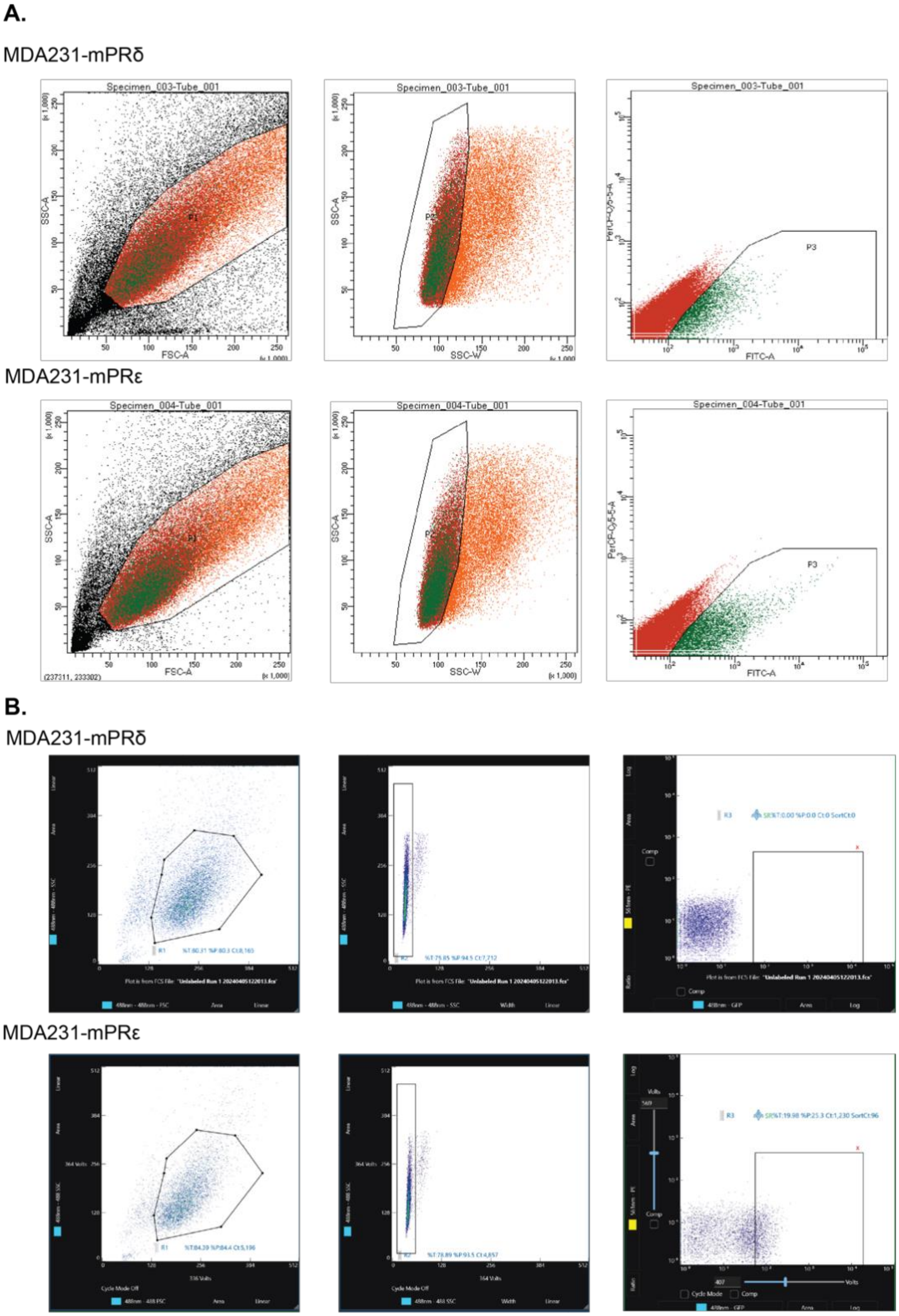

**Supplemental Figure 2. TIRF imaging of mPRδ-GFP and mPRε-GFP cells.** Cultures were fixed, permeabilizied and immunostained with actin together with GFP antibodies and subject to TIRF imaging. Scale bar represents 20μm.

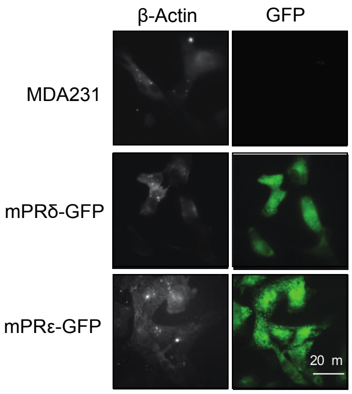


**Supplemental Figure 3. Examining the effects of ORG on PKC activity in cell lines expressing mPRs. A**. Representative immunoblots for MDA231-mPRδ cells treated with ORG (300nM) for 5, 10, 15, or 20 minutes. Cells were lysed, the proteins resolved on SDS-PAGE and subject to immunoblotting Densitometry was carried out to quantify the difference in PKC activation following 300nM ORG treatment after either 5, 10, 15, or 20 minutes. The ratio of phosphorylated kinase to total kinase was determined and normalized to PKC activation at baseline (100%) without ORG treatment (*n*=4).

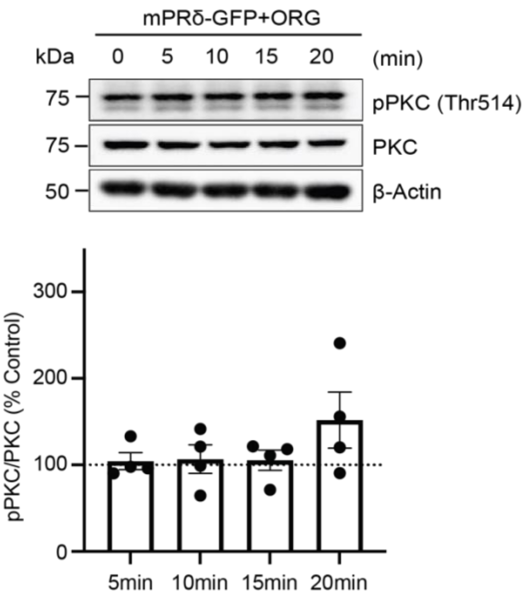

**Supplemental Figure 4: Examining the effects of ORG on protein kinase activity in cell lines. A**. Representative immunoblots for MDA231, MDA231-mPRδ, and MDA231-mPRε cells treated for 20 minutes with increasing concentrations of ORG. Cells were lysed, the proteins resolved on SDS-PAGE, and subject to immunoblotting **B**. Densitometry was carried out to quantify the difference between ORG treatment concentrations and control treatment in MDA231 cells. The ratio of phosphorylated kinase to total kinase was determined and normalized to control treatment (100%) for each cell type (*n*=4). **C**. Densitometry was carried out to quantify the difference between host and mPRδ expressing cells and between ORG treatment concentrations. The ratio of phosphorylated kinase to total kinase was determined and normalized to control treatment (100%) for each cell type (*n*=4). **D**. Densitometry was carried out to quantify the difference between host and mPRε expressing cells and between ORG treatment concentrations. The ratio of phosphorylated kinase to total kinase was determined and normalized to control treatment (100%) for each cell type (*n*=4). Two-Way ANOVA was used to compare treatment responses in MDA231-mPRδ or MDA231-mPRε to MDA231 cells. Post hoc comparisons of each ORG concentration were calculated using the Šidák multiple comparison test.

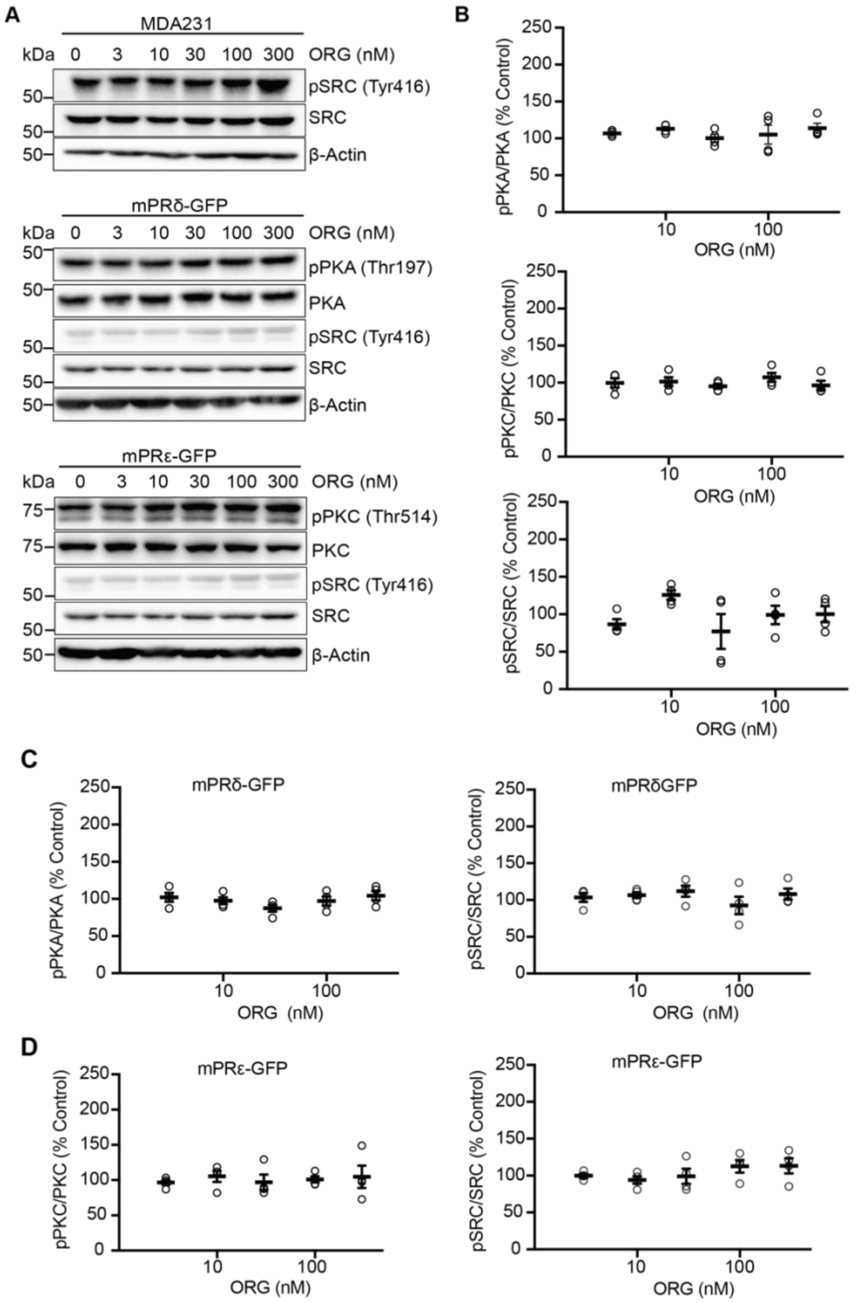

**Supplemental Figure 5. Examining the effects of ORG on protein kinase activity in cell lines. A**. Representative immunoblots for MDA231, MDA231-mPRδ, and MDA231-mPRε cells treated for 20 minutes with ALLO. Cells were lysed, the proteins resolved on SDS-PAGE, and subject to immunoblotting. **B.** Densitometry was carried out to quantify the difference between host and mPR expressing cells and between ALLO treatment concentrations. The ratio of phosphorylated kinase to total kinase was determined and normalized to control treatment (100%) for each cell type (*n*=4). Two-Way ANOVA was used to compare treatment responses in MDA231-mPRδ or MDA231-mPRε to MDA231 cells. Post hoc comparisons of each ALLO concentration were calculated using the Šidák multiple comparison test.

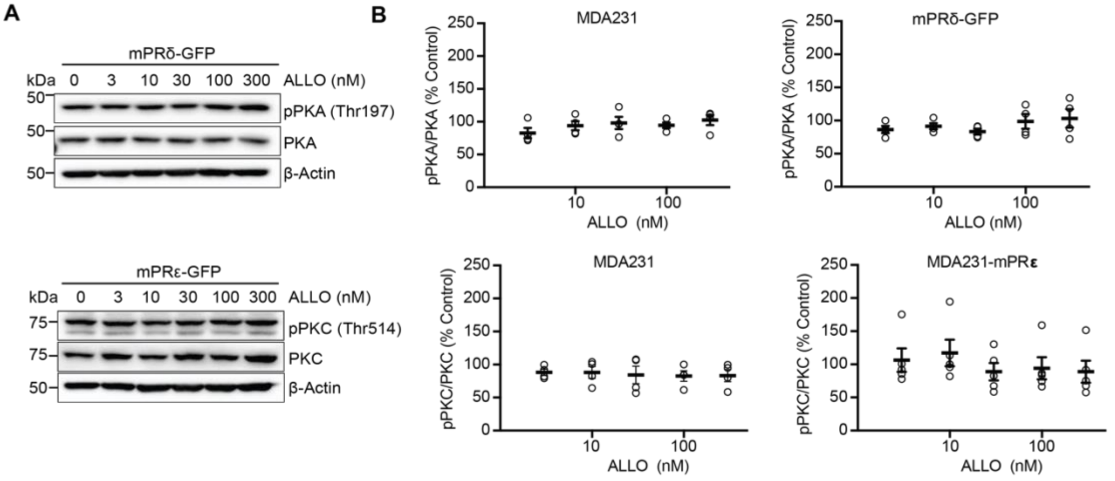


**Supplemental Figure**

**Supplementary Figure 6. Examining the effects of SGE-516 on protein kinase activity in cell lines. A**. Representative immunoblots for MDA231, MDA231-mPRδ, and MDA231-mPRε cells treated for 20 minutes with SGE-516 at 3, 10, 30, 100, and 300nM. Cells were lysed, the proteins resolved on SDS-PAGE, and subject to immunoblotting. **B.** Densitometry was carried out to quantify the difference between host and mPR expressing cells and between SGE-516 treatment concentrations. The ratio of phosphorylated kinase to total kinase was determined and normalized to control treatment (100%) for each cell type (*n*=4). Two-Way ANOVA was used to compare treatment responses in MDA231-mPRδ or MDA231-mPRε to MDA231 cells. Post hoc comparisons of each SGE-516 concentration were calculated using the Šidák multiple comparison test.

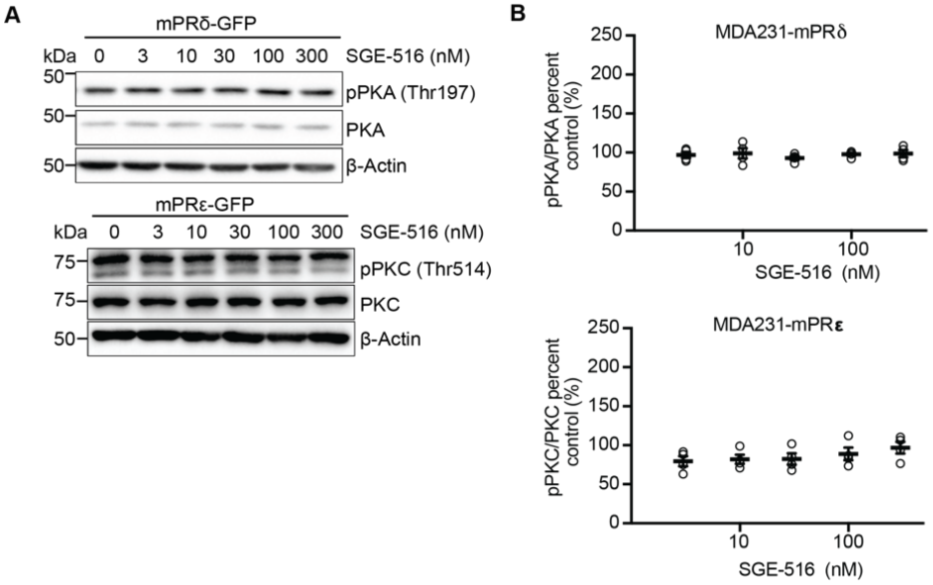


=
